# Hamilton’s rule, gradual evolution, and the optimal (feedback) control of phenotypically plastic traits

**DOI:** 10.1101/2020.09.23.310532

**Authors:** Piret Avila, Tadeas Priklopil, Laurent Lehmann

## Abstract

Most traits expressed by organisms, such as gene expression profiles, developmental trajectories, behavioural sequences and reaction norms are function-valued traits (colloquially “phenotypically plastic traits”), since they vary across an individual’s age and in response to various internal and/or external factors (state variables). Furthermore, most organisms live in populations subject to limited genetic mixing and are thus likely to interact with their relatives. We here formalise selection on genetically determined function-valued traits of individuals interacting in a group-structured population, by deriving the marginal version of Hamilton’s rule for function-valued traits. This rule simultaneously gives a condition for the invasion of an initially rare mutant function-valued trait and its ultimate fixation in the population (invasion thus implies substitution). Hamilton’s rule thus underlies the gradual evolution of function-valued traits and gives rise to necessary first-order conditions for their uninvadability (evolutionary stability). We develop a novel analysis using optimal control theory and differential game theory, to simultaneously characterise and compare the first-order conditions of (i) open-loop traits - functions of time (or age) only, and (ii) closed-loop (state-feedback) traits - functions of both time and state variables. We show that closed-loop traits can be represented as the simpler open-loop traits when individuals do no interact or when they interact with clonal relatives. Our analysis delineates the role of state-dependence and interdependence between individuals for trait evolution, which has implications to both life-history theory and social evolution.

## 1 Introduction

All biological organisms are open systems exchanging energy, matter, and information with their surrounding. As such, most if not all traits of an organism may vary in response to changes of its internal factors as well as to changes in its external biotic and abiotic environmental conditions. Examples include gene expression profiles, physiological processes, reaction norms, life-history traits, developmental trajectories, morphological shapes, and behavioural sequences. We collectively call these traits function-valued traits by which we mean phenotypes whose expression depends on some parameter(s) or variable(s) (e.g., time, space, internal or external biotic and abiotic conditions), and these traits are often called colloquially as “phenotypically plastic traits” in evolutionary biology (West-Eberhard, 2003, p. 33). Formalising how selection shapes these traits is relevant as it helps to understand their evolution and the mechanistic constraints involved in their functioning. This has been done for genetically determined function-valued traits using different theoretical approaches that consider different biological perspectives on the evolution of these traits.

First, the evolution of life-history schedules has often been studied by applying *Pontryagin’s maximum principle* (e.g., León, 1976; Macevicz and Oster, 1976; Oster and Wilson, 1977; Schaffer, 1982; Iwasa and Roughgarden, 1984; Sibly et al., 1985; Stearns, 1992; Perrin, 1992; Kozłowski, 1992; Perrin et al., 1993; Bulmer, 1994; Irie and Iwasa, 2005; Parvinen et al., 2013; Lehmann et al., 2013; Metz et al., 2016). Here, a trait evolves to vary as a function of the age or time of interaction of individuals, while individual fitness (expected survival and reproduction) can be constrained by the dynamics of state variables. These state variables are observables describing internal or external conditions of an individual, e.g., body size, fat reserves, information, resource availability, behaviour of others, that in turn depends on trait expression. Models applying Pontryagin’s maximum principle formalise the evolution of so-called *open-loop* traits (Weber, 2011; Liberzon, 2011), whose name emphasises that the trait itself involves no feedback loop, since it depends only on time (age). As such, an open-loop trait can be thought of as an entirely fixed course of phenotypic expression from birth to death of an individual (trait expression happens “by the clock”); an example would be an age-dependent growth trajectory. Evolution of open-loop traits has also been formalised to include interactions between relatives, which allows to consider their evolution under limited genetic mixing such as spatially or family-structured populations (Bulmer, 1983; Day and Taylor, 1997, 1998, 2000; Wild, 2011; Avila et al., 2019).

Second, in behavioural ecology and evolutionary game theory, selection on function-valued traits has typically been studied by using *dynamic programming* (see Houston et al., 1999; Mangel et al., 1988 for textbook treatments and e.g. Leimar, 1997; Ewald et al., 2007; McNamara and Houston, 1987; Dechaume-Moncharmont et al., 2005). Here, the trait evolves to vary not only as a function of time but also as a function of relevant state variables. This formalises so-called *closed-loop* traits (Weber, 2011; Liberzon, 2011) as these now involve a feedback between trait expression and state dynamics (see Fig. 1 for a schematic conceptualisation). A closed-loop trait can thus be thought of as a contingency plan, which specifies a conditional trait expression rule according to fitness relevant conditions an individual may be in. An example would be an ontogenetic allocation of resources to somatic functions depending on fat reserve (by contrast, the corresponding open-loop trait would allocate resources only depending on age).

**Figure 1:**
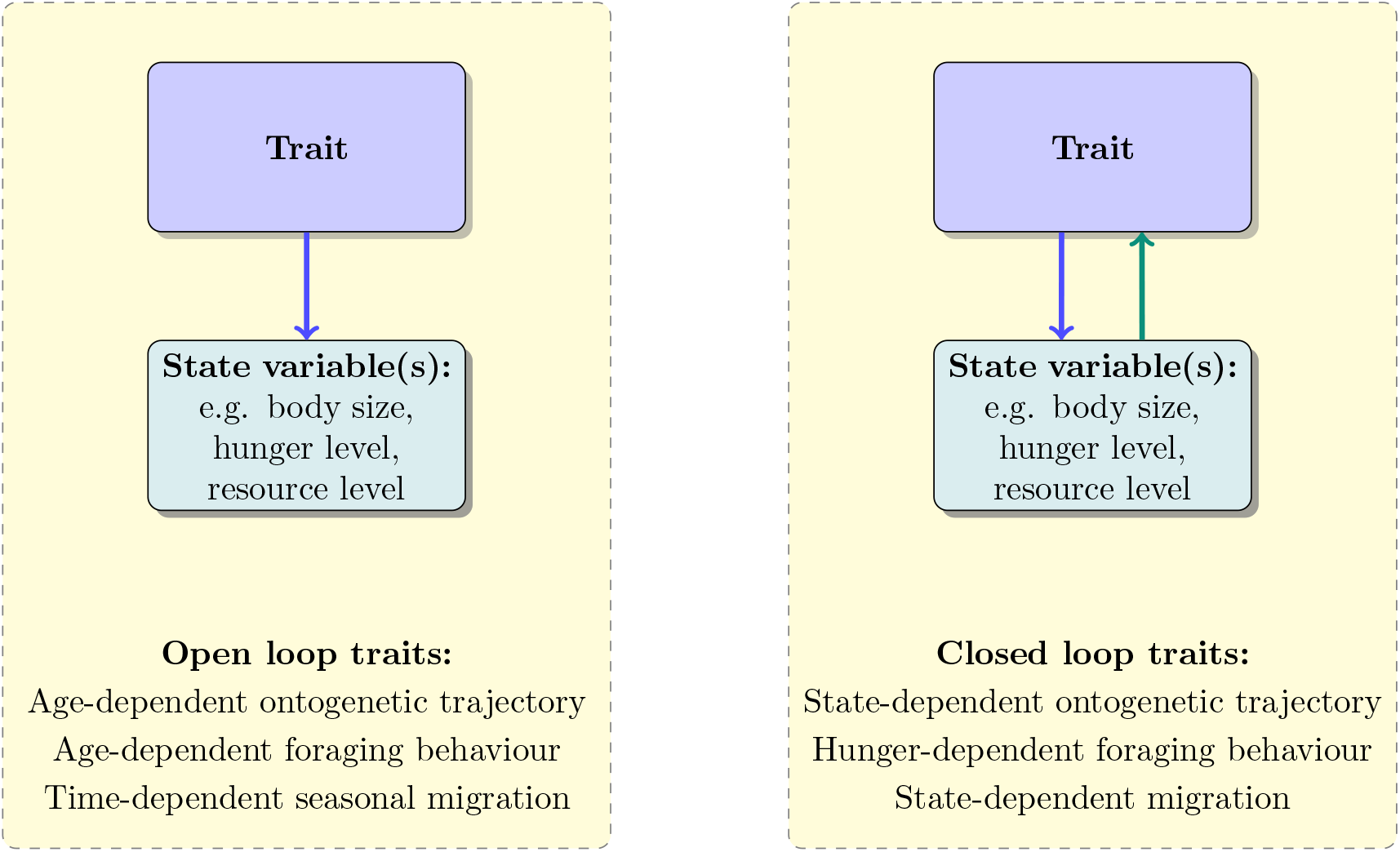
Open-loop (age/time-dependent) and closed-loop (age/time-dependent and state-dependent) conceptualisation of function-valued traits. Open-loop traits affect state variable(s). Closed-loop traits affect state variable(s) and state-variable(s) affect(s) them in turn (thus there is a *feedback-loop*). In both cases: (i) traits and state variables can vary over age/time and (ii) state variables affect fitness. Same biological phenomena can be conceptualised as either open-loop or closed-loop traits.

Both Pontryagin’s maximum principle and dynamic programming are *optimal control theory* approaches (e.g., Bryson and Ho, 1975; Basar and Olsder, 1999; Dockner et al., 2000; Sydsaeter et al., 2005; Weber, 2011; Liberzon, 2011; Kamien and Schwartz, 2012), whose common aim is to identify a schedule of control variables–a trait–over a period of time that maximises (in the best response sense) an objective function (fitness in biology). A crucial result of this literature is that open-loop or closed-loop trait expression leads to different outcomes when individuals interact (e.g. Basar and Olsder, 1999; Dockner et al., 2000). This means that the evolution of a function-valued trait should depend on the assumptions about its functional dependence as well as the type of interactions individuals face. Yet the conditions under which it matters to distinguish between open-and closed-loop traits and how this impacts on an evolutionary analysis remains unclear and has, perhaps surprisingly, not been worked out in evolutionary biology despite the wide interest in the evolution of phenotypic plasticity.

Function-valued traits have also been studied in quantitative genetics theory, where the directional selection coefficient on function-valued traits has been derived assuming no interactions between individuals (Kirkpatrick and Heckman, 1989; Gomulkiewicz and Kirkpatrick, 1992; Gomulkiewicz and Beder, 1996; Beder and Gomulkiewicz, 1998). This selection coefficient describes selection over short timescales (time-scales of demographic changes) and can be decomposed into component-wise descriptions, which allows to describe the direction of selection for each component of a function-valued trait. While this selection coefficient has been connected to long-term evolution and extended to include interactions between individuals in well-mixed populations (Parvinen et al., 2006; Dieckmann et al., 2006), this literature does not distinguish between open-loop and closed-loop traits and it thus remains unclear how the selection coefficient on a trait connects to the dynamics state constraints (that can be physiological or informational) underlying trait evolution. This is why it would be useful to connect the directional selection coefficient on function-valued traits to optimal control theory results, because it provide a way to analyse how selection on traits depends on inter-dependencies and constraints between different trait components.

There are thus different approaches to the evolution of function-valued traits, but the scope of existing results and the connection between them is not clear. In contrast, for selection on quantitative scalar traits general results have long been proven to hold. In particular, for small trait deviations (weak selection), the selection coefficient on a scalar quantitative trait in a population subject to limited genetic mixing can be expressed as a marginal version of Hamilton’s rule, where the direct and indirect fitness effects (the “cost” and “benefit”) are given by partial derivatives of individual fitness (e.g., Taylor and Frank, 1996; Frank, 1998; Roze and Rousset, 2003; Rousset, 2004; Lehmann and Rousset, 2014; Van Cleve, 2015). This selection coefficient provides two useful results about gradual quantitative evolution. First, since the selection coefficient is independent of allele frequency and is of constant sign (Roze and Rousset, 2003, 2004; Rousset, 2004; Lehmann and Rousset, 2014), the marginal Hamilton’s rule subtends gradual evolution of all scalar traits even when the survival and reproduction of individuals depend on the behaviour of others, such as under density- and frequency-dependent selection. Second, when the selection gradient vanishes, Hamilton’s rule provides the necessary first-order condition for a strategy to be locally uninvadable; that is, it allows to determine candidate evolutionary stable strategies, which is central to characterise of long-term evolution (Geritz et al., 1998; Rousset, 2004). Do these general principles of gradual evolution in group-structured populations that hold for scalar traits also hold for function-valued traits?

Our goal in this paper is to formalise the directional selection on genetically determined function-valued traits when state constraints affect trait expression and the evolving population is group-structured (subject to limited genetic mixing). To achieve this we develop a two-step analysis. First, it is to formalise the directional selection coefficient on quantitative function-valued under limited genetic mixing (taking panmictic population as a special case into account) in order to characterise the gradual evolution of function-valued traits. Second, from the the directional selection we derive how state-dependence of trait expression affects selection on function-valued traits. The rest of this paper is organised as follows. (1) We derive the selection coefficient acting on a mutant allele coding for a function-valued trait in the island model of dispersal (group-structured population) under weak selection, which yields the marginal version of Hamilton’s rule for function-valued traits. We deduce from Hamilton’s rule the necessary first-order condition for local uninvadability, which yields the candidate uninvadable function-valued traits and applies to both continuous and discrete traits. (2) We apply these results to time-dependent function-valued traits (dynamic traits) by deriving necessary conditions for uninvadability expressed in terms of dynamic constraints on state variables and their (marginal) effects on the reproductive value. This allows us to compare how selection acts on open-loop versus closed-loop traits, specifying the role of trait responsiveness. In turn, this allows to establish the connection between the dynamic programming and the maximum principle type of approaches in the context of gradual phenotypic evolution. (3) We illustrate the different main concepts of our approach by analysing the evolution of temporal common pool resource production and extraction within groups. (4) Finally, we discuss the scope of our results.

## 2 Model

### 2.1 Biological scenario

Consider a haploid population subdivided into an infinite number of homogeneous groups (without division into class structure) with a fixed number N of individuals, where censusing takes place at discrete demographic time periods. All groups are subject to the same environmental conditions and are equally connected to each other by random dispersal. A discrete demographic time period spans an entire life cycle iteration where various events can occur (e.g. growth, reproduction, dispersal) to individuals. The life cycle may allow for one, several, or all individuals per group to die (thus including whole group extinction through environmental effects or warfare). Generations can thus overlap but when this occurs, the parents are considered equal (in respect to their “demographic properties”) to their offspring in each generation (since there is no within-group class structure). Dispersal can occur before, during, or after reproduction, and more than one offspring from the same natal group can establish in a non-natal group (i.e., propagule dispersal). We refer to this group-structured population where all individuals within groups are indistinguishable, as the *homogeneous island population* (i.e., broadly this corresponds to the infinite island model of dispersal of Wright, 1931, used since at least Eshel, 1972 under various versions to understand selection on social traits, e.g., Rousset, 2004, and where the specifics of our demographic assumptions are equivalent to those considered in Mullon et al. 2016).

We assume that two alleles segregate in the homogeneous island population at a locus of interest: a mutant allele with trait 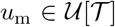 and a resident (wild-type) allele with trait 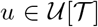. Here, 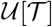 is the set of feasible traits that individuals can potentially express and is formally defined as the set of realvalued functions with range 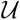 and domain 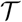, where 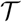 is a space of some index variable(s) representing, for instance, time, an environmental gradient or cue. We assume here that 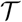 is a closed interval over some discrete or continuous index variable *t*. If *t* is discrete, then the element 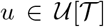 is a vector and if *t* is continuous then the element 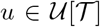 is a piece-wise continuous function. Hence, we write 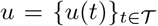 to emphasise that it consists of the (finite or infinite) collection of all point-wise values *u*(*t*) of the trait. Namely, *u* can be thought of as a continuous or a discrete “path” (or a schedule, or a trajectory) on the space 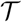. Note that in Table 1 we have outlined a list of symbols of key components of our model.

**Table 1:** Symbols of key components of the model

|  |  |  |
| --- | --- | --- |
| Control variables<br>(evolving traits) | $u, u_m$ | resident and mutant trait |
| | $u^*$ | candidate uninvadable trait |
| | $u_\bullet, u_o, u$ | focal's trait, average neighbour's trait, and average trait of individuals in other groups |
| | $\mathbf{u}_\bullet = (u_\bullet, u_o, u)$ | a vector collecting traits from the focal's perspective |
| | $u = \{u(t)\}_{t \in \mathcal{T}} \in \mathcal{U}[\mathcal{T}]$ | all traits considered here (except section 2.3.1. where we consider scalar traits) belong to a set of real-valued function with range $\mathcal{U}$ and domain $\mathcal{T}$ |
| Decision rules | $d(t, \mathbf{x}(t))$ | closed-loop (feedback) control |
| | $d(t)$ | open-loop control |
| | $d(\mathbf{x}(t))$ | stationary (closed-loop) control |
| Components<br>describing<br>fitness | $w(u_\bullet, u_o, u)$ | focal individual's fitness |
| | $f(t, \mathbf{u}_\bullet(t), \mathbf{x}_\bullet(t))$ | rate of fitness increase at time $t$ |
| | $\Phi(\mathbf{x}_\bullet(t_f))$ | fitness increase at final time $t_f$ (scrap value) |
| | $v(t, \mathbf{x}(t); u)$ | (neutral) reproductive value function |
| | $\boldsymbol{\lambda}(t, \mathbf{x}(t); u)$ | a vector of shadow values of state variables ($\boldsymbol{\lambda}(\cdot) = (\lambda_\bullet(\cdot), \lambda_o(\cdot), \lambda(\cdot))$); the shadow value of a state variable gives the future fitness effect from marginally changing the state ($\boldsymbol{\lambda}(t, \mathbf{x}(t); u) = \nabla v(t, \mathbf{x}(t); u)$) |
| | $H$ | Hamiltonian function, contribution to individual fitness at time $t$ due to current “activities” |
| Components<br>describing<br>selection | $s_\eta(u)$ | selection coefficient |
| | $s(u)$ | selection gradient ($s_\eta(u) = \epsilon \eta \cdot s(u)$) |
| | $\epsilon$ | effect size of the mutant allele (scalar) |
| | $\eta$ | phenotypic deviation of the mutant allele (function; $\eta = \{\eta(t)\}_{t \in \mathcal{T}}$) |
| | $-c_\eta(u), b_\eta(u)$ | direct and indirect fitness effects |
| | $-c(t, u), b(t, u)$ | point-wise direct and indirect fitness effects |
| | $r(u)$ | (neutral) relatedness between two randomly sampled (without replacement) group members |
| State variables | $x, x_m$ | resident and mutant state |
| | $x^*$ | candidate uninvadable state |
| | $x_\bullet, x_o, x$ | focal's state, average neighbour's state, and average state of individuals in other groups |
| | $x = \{x(t)\}_{t \in \mathcal{T}}$ | all state variables $x(t) \in \mathbb{R}$ considered are real-valued functions defined over domain $\mathcal{T}$ |
| | $g(\mathbf{u}_\bullet(t), \mathbf{x}_\bullet(t))$ | function describing the rate of change of a state variable (here for the focal individual) |
| | $\mathbf{g}(\mathbf{u}_\bullet(t), \mathbf{x}_\bullet(t))$ | A vector collecting the change rates in state variables of the representative individuals, i.e. $\mathbf{g}(\mathbf{u}_\bullet(t), \mathbf{x}_\bullet(t)) = (g(\mathbf{u}_\bullet(t), \mathbf{x}_\bullet(t)), g(\mathbf{u}_o(t), \mathbf{x}_o(t)), g(\mathbf{u}(t), \mathbf{x}(t)))$ |

The crucial assumption of this paper is that the mutant trait *u*_m_ can be expressed in the following form as a small deviation from the resident trait:

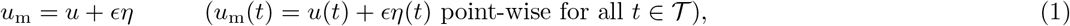

where the phenotypic deviation function 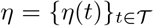 must satisfy 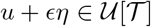 for sufficiently small non-negative parameter *ϵ*. Because 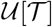 may have a boundary, not all phenotypic deviations generate a mutant strategy *u*_m_ that remains within the bounds of the feasible trait space 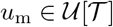, independent of the choice of e (see Section 2.3.2). Note that we are making a distinction between a phenotypic deviation *η* (a function) and the effect size e (a scalar) of that deviation. In the literature, a (scalar) mutant effect is often modelled with the notation *δ* = *ηϵ* (e.g. Rousset, 2004). This distinction between phenotypic deviation and effect size in the notation is necessary for analysing selection on function-valued traits.

### 2.2 Allele frequency change and short-term evolution

Our first aim is to characterise the change in mutant allele frequency in the homogeneous island population under weak selection (*ϵ* ≪ 1). To that end, it is useful to follow the direct fitness approach (Taylor and Frank, 1996; Rousset and Billiard, 2000; Rousset, 2004) and introduce the individual fitness function 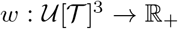 such that *w*(*u*_•_, *u*_o_, *u*) gives the expected number of successful offspring produced over one life cycle iteration by a focal individual (possibly including self through survival) with trait u_•_, when its average neighbour in the focal group has trait uo and an average individual (from other groups) in the population has trait u, which is taken here to be the resident trait for simplicity of presentation. We note that any individual in the population can be taken to be the focal individual (Rousset and Billiard, 2000; Rousset, 2004) and that the fitness of this individual can always be expressed in terms of average phenotypes of other individuals in different roles with respect to the focal (e.g., group neighbour, cousin, members of other groups, etc.), whenever mutant and resident phenotypes are closely similar (see the argument in Appendix A.2 for function-valued traits and a textbook argument for scalar traits e.g. Rousset, 2004, p. 95). These individuals in different roles, as well as the focal individual itself, are actors on the fitness of the focal. Here, the focal individual is regarded as the recipient of the trait expressions of different actors (i.e. focal individual, average neighbour, average individual in other groups), which corresponds to the direct fitness or recipient-centred approach (e.g. Rousset, 2004, Chapter 7).

In terms of this definition of individual fitness, we define the *direct fitness effect* of expressing the mutant allele as

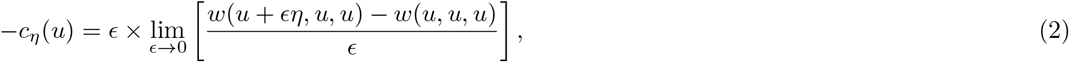

which is the effect that the focal individual has on its own fitness if it would switch from expressing the resident to the mutant allele for a small allelic effect. Analogously, we define the *indirect fitness effect* of expressing the mutant allele as

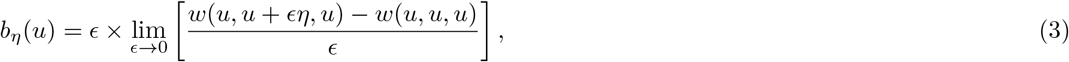

which is the effect that the whole set of neighbours have on focal’s fitness if they were to all switch from expressing the resident to the mutant allele (note that e appears in front of the derivatives in eq. (2)–(3) because it scales the fitness effect of a trait deviation in direction *η* and so –*c_η_*(*u*) and *b_η_*(*u*) are the net fitness effects). Finally, let us denote by *r*(*u*) the neutral relatedness between two randomly sampled group neighbours (Michod and Hamilton, 1980; Frank, 1998; Rousset, 2004) in the homogeneous island population that is monomorphic for the resident; namely, r(u) is the probability that in a neutral process (where all individuals are alike) the two homologous alleles of these individuals coalesce in the same common ancestor (e.g., Roze and Rousset, 2003; Rousset, 2004; Lehmann and Rousset, 2014; Van Cleve, 2015). Note that relatedness defined as such depends only on the resident trait. In Appendix A, we show that the change Δ*p* in the frequency *p* of the mutant allele over one demographic time period (one life cycle iteration) can be expressed in terms of these quantities as follows.

#### Invasion implies substitution principle result

*In the homogeneous island population with two alleles, the change in mutant allele frequency p in the population takes the form*

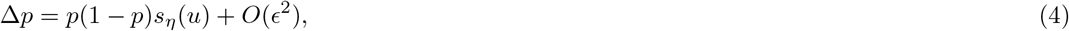

*where p*(1 – *p*) *is the genetic variance at the locus of interest*,

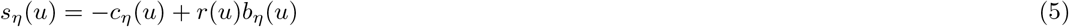

*is a selection coefficient of order O*(*ϵ*)*that is independent of p, and O*(*ϵ*^2^) *is a remainder of all higher order terms. This entails an “invasion implies substitution” property of the mutant allele, which says that if s_η_* (*u*) *> 0, the mutant allele coding for a small function-valued deviation ϵη is selected for and not only invades but substitutes the* (*ancestral*) *resident allele [since effects of order O*(*ϵ*^2^) *can be neglected in* *eq.* (4) *whenever s_η_* (*u*) *is non-zero]*.

We have thus formalised an “invasion implies substitution”-principle (see Priklopil and Lehmann, 2020 for a review) for function-valued traits in the homogeneous island population and which takes the form of Hamilton’s rule: the mutant spreads if *r*(*u*)*b_η_*(*u*) – *c_η_* (*u*) > 0. This novel result is a multidimensional generalisation of previous analogous results for scalar traits (Roze and Rousset, 2003, 2004; Rousset, 2004; Lehmann and Rousset, 2014).

Owing to its simplicity, the function-valued trait nature of our result is perhaps yet not fully apparent, but is made explicit on noting that the direct and indirect effects (eqs. 2–3) are both formally *Gâteaux derivatives*, which are directional derivatives (see section A.1 in Appendix for a formal definition and e.g., Troutman, 1991, p. 45-50, Luenberger, 1997, p. 171-178) and represent change in fitness resulting from a sum of all weighted component-wise changes in trait expression (over the domain 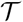) induced by the mutation function *η*. To outline the component-wise change in fitness, it is useful to decompose the selection coefficient as

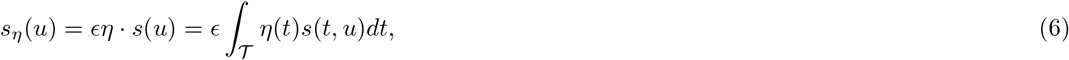

where o is an inner product on functions (the generalisation of a dot product, see e.g. Anton and Rorres, 2013, Chapter 6), 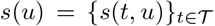 is the selection gradient function, where the component *s*(*t*,*u*) gives the selection gradient on component *u*(*t*) of the trait, i.e. the value of *u* at time *t*, holding other components *u*(*t′*) (for all 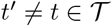) of the trait fixed. Each component of the selection gradient function is then given by

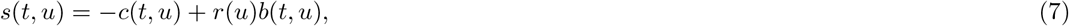

where

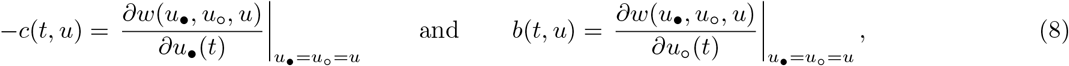

are, respectively, the effect on the focal’s own fitness from changing marginally component *u*_•_(*t*) of its trait, while holding other trait components *u*_•_(*t′*) (for *t′ ≠ t*) fixed, while *b*(*t*, *u*) is the effect of all group neighbours on the focal individuals fitness when changing marginally component *u*_o_(*t*) of their traits, while holding other components *u_o_*(*t′*) (for *t′ ≠ t*) of their traits fixed. That is, the costs and benefits are partial derivatives and *s*(*t*, *u*) is the inclusive fitness effect on a focal individual. When *t* is discrete and 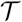 finite, eq. (7) corresponds to the trait specific inclusive fitness effect derived previously for a backdrop monomorphic resident homogeneous island population (Mullon et al., 2016, eq. 12).

Eq. (6) shows that the selection coefficient is a weighted change of trait-specific changes. Note that for continuous index variable *t* over the interval 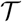, the partial derivatives *–c*(*t*, *u*) and *b*(*t*, *u*) in eq. (8) are formally functional derivatives (e.g. Troutman, 1991, p. 45-50, Luenberger, 1997, p. 171-178). In the absence of interactions between relatives *s*(*t*,*u*) reduces to *β*(*y*) in eq. 1 of Gomulkiewicz and Kirkpatrick (1992) for *y = t*, *G*(*a*) in eq. 4 of Parvinen et al. (2006) for *a* = *t*, or g(*a*; *u*) in eq. 3 in Metz et al. (2016) for *a* = *t* (but see Δ*W*_incl_(*t*) in eq. 25 of Day and Taylor, 2000, which allows for interactions between relatives).

### 2.3 Necessary condition for local uninvadability and long-term evolution

It follows from our “Invasion implies substitution principle” result that a necessary first-order condition for a trait *u^*^* to be locally uninvadable (resistant to invasion by any mutant in a small radius *ϵ* ≪ 1) is given by a non-positive selection coefficient for all admissible mutants in the resident *u^*^* population, that is

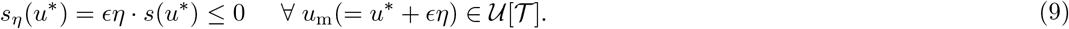

Local resistance to invasion by sets of alternative mutants allows to characterise candidate long-term evolutionary outcomes (Eshel and Feldman, 1984; Eshel, 1996; Eshel et al., 1998) and is a first-step (and often the only accessible computational step under limited genetic mixing) towards characterising uninvadable traits.

A crucial question is whether a locally uninvadable strategy *u^*^* will be approached by gradual evolution from within its neighbourhood and thus be convergence stable (Eshel, 1983; Lessard, 1990; Geritz et al., 1998; Rousset, 2004; Leimar, 2009). Because characterising convergence stability involves a second order analysis of the selection coefficient, which is involved for multidimensional traits (Lessard, 1990; Leimar, 2009), it will not be investigated further in this paper. For the same reason, we will also not consider sufficient conditions for local uninvadability. In the remainder of this section, we focus on characterising in more detail the necessary condition of local univadability (eq. 9) in terms of the selection gradient function *s*(*u*), which allows removing the considerations of mutational effect *η*.

#### 2.3.1 Local uninvadability for scalar-valued traits

Let us first consider the case of scalar quantitative traits, where the trait of each individual is an element belonging to a bounded subset 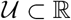 of the real line. That is, the resident and mutant traits stay within the feasibility bounds *u*_min_ ≤ *u, u*_m_ ≤ *u*_max_). For this case, the index *t* in eq. (7) can be dropped and one obtains the standard selection gradient on a scalar-valued trait for the homogeneous island population:

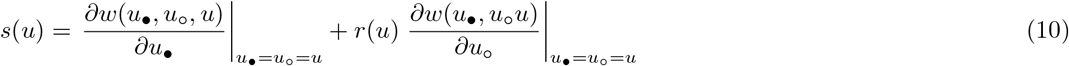

(Taylor and Frank, 1996; Frank, 1998; Roze and Rousset, 2003, 2004; Rousset, 2004; Lehmann and Rousset, 2014; Van Cleve, 2015).

Note that for *u* = *u*_min_ an admissible phenotypic deviation *η* must be non-negative *η* ≥ 0 and for *u* = *u*_max_ it must be non-positive *η* ≤ 0 while for *u*_min_ ≤ *u* ≤ *u*_max_ the deviation *η* is unrestricted. Substituting this into the first-order condition for uninvadability eq. (9) yields that the necessary condition for uninvadability for scalar bounded traits can be expressed in the following form

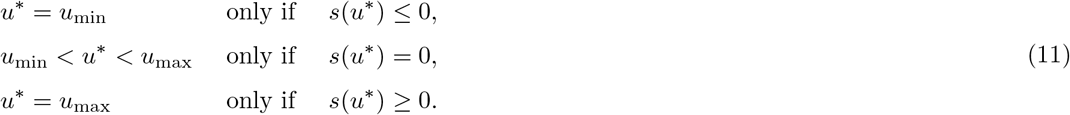

Note that if the set of admissible traits is unbounded (i.e. 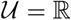), then the first-order necessary condition for local uninvadability is given by the second line of eq. (11).

#### 2.3.2 Local uninvadability for function-valued traits

Let us return to the general case where the trait of each individual is an element of 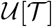, being either a vector (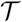 discrete) or a (bounded and piece-wise continuous) function (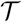 continuous). More precisely, for all 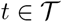 the resident and mutant traits stay within the feasibility bounds *u*_min_(*t*) ≤ *u*(*t*), *u*_m_(*t*) ≤ *u*_max_(*t*), such that 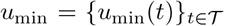 and 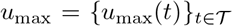. Now, an admissible deviation 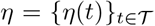 must satisfy for all 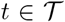 similar conditions as given for the scalar-traits in Section (2.3.1), that is, for *u*(*t*) = *u*_min_(*t*) an admissible phenotypic deviation *η* must be non-negative *η*(*t*) ≥ 0 and for *u*(*t*) = *u*_max_(*t*) it must be non-positive *η*(*t*) ≤ 0 while for *u*_min_(*t*) < *u*(*t*) < *u*_max_(*t*) the deviation *η*(*t*) is unrestricted. Substituting the admissible deviations into eq. (9) yields that a candidate uninvadable strategy 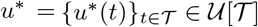 satisfies for all 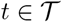:

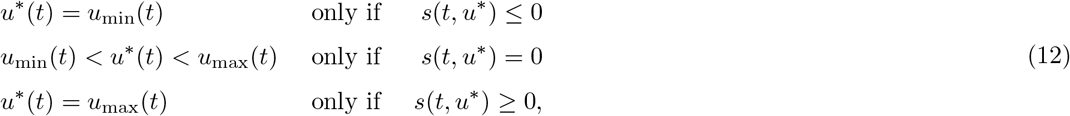

which is thus equivalent to eq. (11) in a point-wise way.

## 3 From the selection gradient to candidate optimal controls

The point-wise description of the candidate uninvadable trait *u^*^* given by eq. (12) is unlikely to be directly useful in solving for *u^*^* in concrete applications because factors characterising the organism and its environment change over time (e.g. organisms can grow and the resources in the environment can get depleted). Hence, solving *u^*^* from eq. (12) would entail simultaneously solving a large number of equations, while tracking the changes in the relevant time-dependent factors. A more useful characterisation of *u^*^* can be achieved with the use of the mathematical framework of optimal control theory, most notably dynamic programming and Pontryagin’s maximum principle, both of which have been used abundantly in evolutionary biology and in different contexts (e.g., León, 1976; Iwasa and Roughgarden, 1984; Mangel et al., 1988; Houston et al., 1999; Stearns, 1992; Perrin, 1992; Perrin et al., 1993; Kozłowski, 1992; Day and Taylor, 1997, 2000; Cichon and Kozlowski, 2000; Irie and Iwasa, 2005; Lehmann et al., 2013; Priklopil et al., 2015; English et al., 2016; Metz et al., 2016; Avila et al., 2019).

## 3.1 Key concepts

### 3.1.1 Fitness function, control variables and state constraints

For space reasons, we focus on a continuous time formulation (but parallel developments apply to discrete time), and assume that a demographic time period is characterised by the time interval 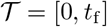 during which the trait expression is observed. This time interval can be thought of as the length of the lifespan of organisms or the time during which behavioural interactions occur between individuals (e.g. a mating season, winter season), which eventually leads to reproduction. More specifically, we now assume that the fitness of the focal individual can be written in the form

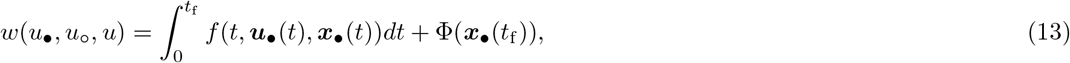

where *f* (*t*, ***u***_•_(*t*), ***x***_•_(*t*)) is the rate of increase of individual fitness at time *t* and Φ(***x***_•_(*t_f_*)) is the so-called scrap value; namely, the contribution to individual fitness at the final time *t* = *t*_f_ (formally 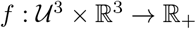 and 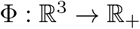). Here,

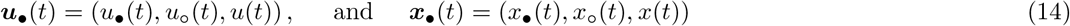

collect, respectively, the trait expression levels *u*_•_(*t*), *u_o_*(*t*), and *u*(*t*) at time t of the focal individual, that of an average neighbour, and an average individual from the population, and the *state variables x*_•_(*t*), *x_o_*(*t*), and *x*(*t*) of these respective individuals (note that the “•” in the subscript of ***u***_•_ and ***x***_•_ emphasises that these controls and state variables collect those of all actors on the fitness of the focal recipient). State variables describe some measurable conditions of an individual (e.g. size, stored energy, hunting skill) or that of its environment (e.g. amount of resource patches, environmental toxicity). The defining feature of a state variable in our model is that its time dynamics depends on the evolving trait of one or more individuals in interaction and we will henceforth from now on call the elements of ***u***_•_(*t*) the *control variables*, which is customary for these type of models in the evolutionary literature (e.g., Perrin, 1992; Day and Taylor, 2000).

Because models with both control and state variables become rapidly notationally complex, we assume that both the controls and the state variables are one-dimensional real numbers. The state of every individual is assumed to change according to the function 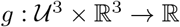, such that

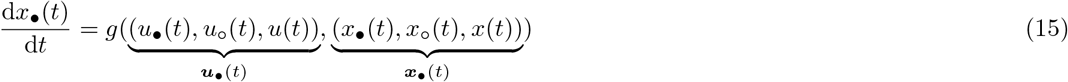

with initial condition (“i.c.”) *x*_•_(0) = *x*_•_ = *x*_init_ and which is the rate of change of the state of a focal individual with control *u*_•_(*t*) in state *x*_•_(*t*), when its neighbours have average control *u*_o_(*t*) and average state *x*_o_(*t*) in a population where the average control (in other groups) is *u*(*t*) and the average state is *x*(*t*). Similarly, we can also express the rate of change of the state of an average neighbour of the focal and an average individual in the rest of the population, respectively, as

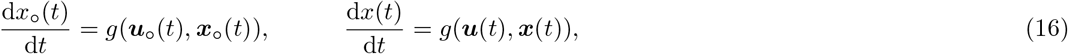

where the vectors

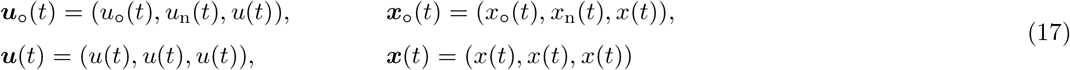

collect the (average) controls and states of actors on the state variables of an average neighbour of the focal individual (first line), and on an average individual in the population (second line), respectively (here and throughout all vectors are defined by default as being column vectors). These actors are thus second-order level actors on the focal recipient since they affect the state variables of actors affecting the focal’s fitness. Note that the subscripts of the control vectors (***u***_o_(*t*) and ***x***_o_(*t*)) and state vectors (***u***(*t*) and ***x***(*t*)) emphasise the individual (actor) from who’s perspective the second-order actors’ control and variables are collected. Accordingly, the vectors in eq. (17) contain elements

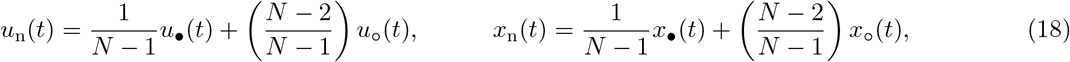

which are, for an average neighbour of the focal, the control and state expressions of average neighbours viewed as actors on the focal individual. While we have so far explicitly distinguished between the states of different individuals, which is required if state represents some property of individual’s condition (e.g. body size or individual knowledge), nothing prevents the individual state to represent some environmental condition common to the group or population and which can be influenced by individual behaviour (e.g. local amount of resources in the group, in which case *x*_•_(*t*) = *x*_o_(*t*), see concrete example in section 4). Note that while tracking the dynamics of three state variables (eqs. 15–18) may appear complicated, it is much simpler than tracking the state of all individuals in a group separately (which would require as many equations as there are individuals in the group and is the approach taken in Day and Taylor, 1997, 2000 and differential game theory, e.g., Dockner et al., 2000; Weber, 2011).

Finally, we now make a couple of remarks about the properties of the fitness function *w*(*u*_•_, *u*_o_, *u*) (eq. 13) and its dynamic constraints (eqs. 15–16), which is a special case of a fitness function *w*(*u*_•_, *u*_o_, *u*) considered in section 2. First, the fitness function (13) depends on the full trajectories of the control 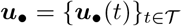 and state 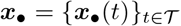 variables, but since the state variables are fully determined by the controls (by way of eqs. 15–16) and the initial condition x_init_ (which we assume here to be fixed), then fitness is determined by the controls. In particular, if fitness depends only on the state of the system at the final time *t*_f_ (*w*(*u*_•_, *u*_o_, *u*) = Φ(***x***_•_ (*t*_f_))), then fitness still depends critically on the control variables. We assumed in section 2 that the fitness *w*(*u*_•_, *u*_o_, *u*) is Gâteaux differentiable (eqs. 2 and 3), which means here that functions *f*, Φ and *g* are smooth enough with respect to its arguments (see e.g. section 3 of Liberzon, 2011 for textbook treatment of assumptions and Clarke, 1976 for minimal assumptions needed). We finally note that in the homogeneous island population, individual fitness depends in a non-linear way on various vital rates (e.g, Roze and Rousset, 2003, eq. 35, Akçay and Van Cleve, 2012, eq. A12, Van Cleve, 2015, eq. 38, Mullon et al., 2016, eq. box 1a), which themselves may depend on integrals depending on the control schedules of the individuals in interaction. Such situations can be analysed either by defining state variables whose integrated values represent the integral, and are covered by the scrap value Φ(***x***_•_(*t*_f_)) in eq. (13), or by noting that to the first-order, functions of integrals can be replaced by integrals of first-order Taylor series of fitness and hence the *f*(***u***_•_(*t*), ***x***_•_(*t*)) fitness component in eq. (13) may be evaluated as a first-order Taylor expansion of fitness in its vital rates (e.g., Van Cleve, 2015, eq. 39, Mullon et al., 2016, eq. A60–A61).

### 3.1.2 Concept of neutral reproductive value and shadow value

A central role in our analysis will be played by the neutral future-value reproductive value

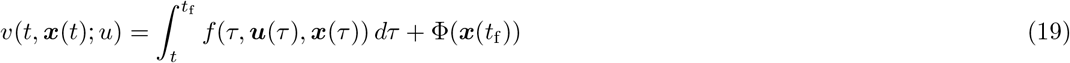

of an individual at time t in a resident population, which gives the total contribution to fitness from time *t* onward of a (recipient) individual when the current state variables of the actors on its fitness is ***x***(*t*). We emphasise that the future-value reproductive value or residual fitness *v*(*t*, ***x***(*t*); *u*) is connected but not equivalent to Fisher’s reproductive value (e.g., Goodman, 1982; Charlesworth, 1994), which is the residual fitness conditional on reaching a certain age (a current-value reproductive value). The argument u has been separated with the semicolon in order to emphasise that the (future-value) reproductive value *v*(*t*, ***x***(*t*); *u*) is evaluated assuming a fixed control trajectory where u is treated as a parameter. Hence, the reproductive value is formally a function of current time *t* and state 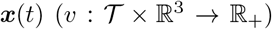. In Appendix B.1, we show that the reproductive value satisfies the following partial differential equation (PDE)

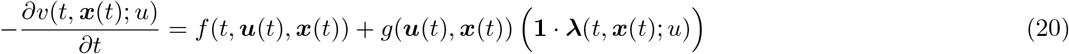

with final condition (“f.c.”) *v*(*t*_f_, ***x***(*t*_f_); *u*) = Φ(***x***(*t*_f_)), where **1** = (1,1,1), “·” is the inner product of vectors and the vector

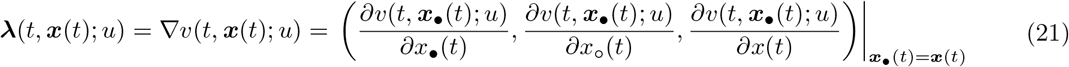

is the gradient of the reproductive value with respect to the changes in the state variables of each individual affecting the focal’s fitness (and associated with fixed resident control path *u*). In the last equality, we use the vector ***x***_•_(*t*) as an argument of the reproductive value (which is defined in a resident population), which might be confusing at first glance, since all individuals in the resident population are actually in the same state. However, the reason why we use the vector ***x***_•_(*t*) when expressing the partial derivatives is because we want to emphasise which state (focal indidivual’s, average neighbour’s or average in other groups) we are varying.

The “-” sign on the left-hand-side of eq. (20) indicates that the reproductive value of an individual is growing when looking backwards in time. Hence, it grows according to the current rate *f*(*t*, ***u***(t), ***x***(*t*)) of fitness increase and the sum **1** · **λ**(*t*, ***x***(*t*); *u*) of the effects of the current state change of each type of actor on the future fitness of the focal individual, weighted by the change g(***u***(*t*), ***x***(*t*)) of state of the actors that are all the same in a resident population. The elements of the gradient **λ**(*t*, ***x***(*t*); *u*) are called the *shadow values* of the states in the optimal control literature (see e.g. Dorfman, 1969; Caputo and Caputo, 2005), since by changing state, there is no immediate effect on fitness, but only future fitness effects.

### 3.1.3 Concept of open and closed-loop controls

Because the internal and external conditions of organisms vary, trait expression can evolve to be functionally dependent on these conditions (Sibly and McFarland, 1976; McFarland, 1977; McFarland and Houston, 1981; Houston et al., 1999). Hence, trait expression can depend on time and state variables. Focusing on the resident trait *u*(*t*), we can conceptualise trait expression in (at least) two different ways that are relevant to evolutionary biology. Namely,

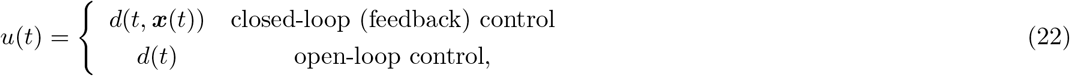

where the function 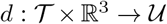 is the trait expression rule (or *decision rule* for short) in the so-called *closed-loop* (or feedback or Markovian) form of the control variable (Basar and Olsder, 1999, p. 221, Dockner et al., 2000, p. 59).

### 3.2 First-order conditions for closed-loop controls

#### 3.2.1 The first-order condition in terms of dynamic constraints

Let us now evaluate the point-wise fitness effects of Hamilton’s marginal rule (7) by substituting the fitness function eq. (13) into eq. (8) and taking the derivative with respect to *u*_•_(*t*) and *u*_o_(*t*). Calculations displayed in Appendix B (in particular eqs. B.16–B.28) then show that the direct effect is

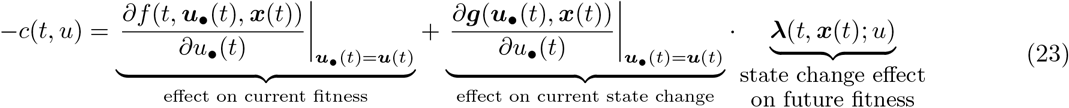

and the indirect effect is

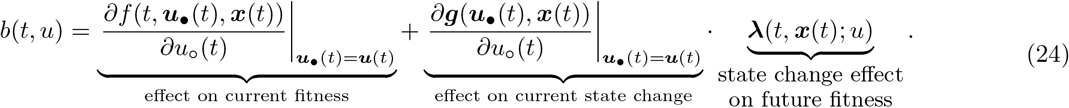

The derivatives are evaluated at 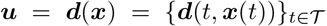 for closed-loop controls and at 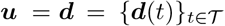 for open-loop controls, where ***d***(*t*, ***x***(*t*)) = (*d*(*t*, ***x***(*t*)), *d*(*t*, ***x***(*t*)), *d*(*t*, ***x***(*t*))) and ***d***(*t*) = (*d*(*t*), *d*(*t*), *d*(*t*)) are vectors of closed-loop and open-loop trait expression rules, respectively, evaluated in a resident population.

We now make two observations about the direct and indirect effects. First, the perturbations of the change of the state variables (*∂****g***(***u***_•_(*t*), ***x***(*t*))/*∂****u***_•_(*t*) and *∂****g***(***u***_•_(*t*), ***x***(*t*))/*∂u*_o_(*t*)) have cascading downstream effects on fitness growth rate *f* (*t*, ***u***_•_(*t*), ***x***_•_(*t*)), but since under a first-order analysis everything else than the original perturbation needs to be held constant, the downstream effects are accounted for by the shadow values **λ**(*t*, ***x***(*t*); *u*) evaluated in the resident population. Second, the state dynamics of an average individual in the population is not affected from variations in *u*_•_ and *u*_o_ (*∂g*(***u***(*t*), ***x***(*t*))/∂u_•_(*t*) = *∂g*(***u***(*t*), ***x***(*t*))/*∂u_o_*(*t*) = 0, by way of eq. 16).

In order to obtain a full characterisation of the first-order condition taking into account the dynamic constraints brought by the shadow value, it is useful to introduce the Hamiltonian function

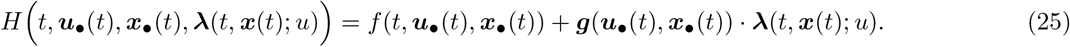

This can be thought of as the contribution to individual fitness of all current “activities” (Dorfman, 1969, p. 822); namely, the (phenotypic) expressions ***u***_•_(*t*) of all individuals currently in state ***x***_•_(*t*) at *t*, holding everything else fixed in a resident population. It is thus the sum of the current rate of fitness contribution *f* (*t*, ***u***_•_(*t*), ***x***_•_(*t*)) and the changes in states ***g***(***u***_•_(*t*), ***x***_•_(*t*)) = (*g*(***u***_•_(*t*), ***x***_•_(*t*)), *g*(***u***_o_(*t*), ***x***_o_(*t*)), *g*(***u***(*t*), ***x***(*t*))) weighted by **λ**(*t*, ***x***(*t*); *u*) evaluated in the resident population, since the shadow values do not directly depend on the activities at time *t*. Our next result (proved in Appendices B.1 and B.2) establishes the necessary condition for uninvadability for closed-loop control paths as follows.

#### Closed-loop control result

*Let 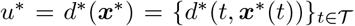 *be a candidate uninvadable closed-loop control path with associated state path* 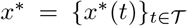, where* ***x***^*^(*t*) = (*x*^*^(*t*), *x*^*^(*t*), *x*^*^(*t*)) *and shadow value* **λ**^*^(*t*, ***x***^*^(*t*)) = **λ**(*t*, ***x***^*^(*t*); *u*^*^). *The candidate uninvadable control path d*^*^(***x***^*^) *has to necessarily satisfy* *eq.* (12), *where the point-wise selection coefficient s*(*t*,*u*^*^) *on control component u*^*^(*t*) = *d*^*^(*t*, *x*^*^(*t*)) *can be written for all* 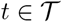 *as*

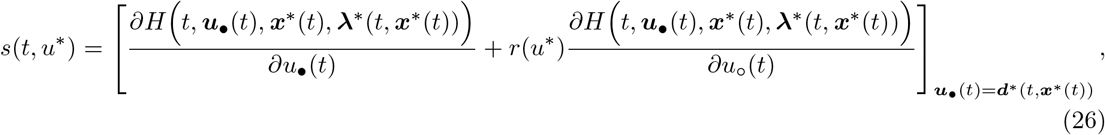

*where* ***d***^*^(*t*, ***x***^*^(*t*)) = (*d*^*^(*t*, ***x***^*^(*t*)), *d*^*^(*t*, ***x***^*^(*t*)), *d*^*^(*t*, ***x***^*^(*t*))), *the state variable satisfies*

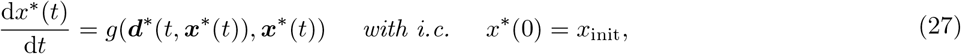

*and the shadow value function* ***λ***^*^(*t*, ***x***^*^(*t*)) = ∇*v*^*^(*t*, ***x***^*^(*t*)) *is obtained from the reproductive value function v*^*^(*t*, ***x***^*^(*t*)) = *v*(*t*, ***x***^*^(*t*); *u*^*^) *that satisfyies*

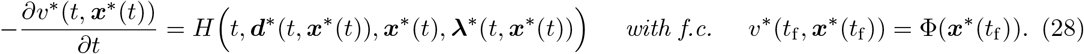

We now emphasise two points about this result where all quantities are evaluated on the resident control *u*^*^ = *d*^*^(*x*^*^) and state *x*^*^ paths. First, the dynamic constraints entail solving forward in time eq. (27), which is an ODE (ordinary differential equation), and solving backwards in time eq. (28), which is a PDE (partial differential equation). Thus, the Hamiltonian can be thought as the growth rate of the reproductive value (when looking backwards in time). For a reader familiar with the dynamic programming literature, the reproductive value *v*^*^(*t*, ***x***^*^(*t*)) is not the so-called *value function* of the model and hence eq. (20) (even when evaluated along the candidate uninvadable control path ***u***_•_ = ***u***^*^) is not the eponymous *Hamilton-Jacobi-Bellman* equation (e.g., Bryson and Ho, 1975; Kamien and Schwartz, 2012; Basar and Olsder, 1999; Dockner et al., 2000; Liberzon, 2011; Weber, 2011). This means that the above result says nothing about the sufficiency of uninvadability, like any standard first-order selection analysis. In this regard, our result provides a weaker, yet simpler and novel condition to characterise closed-loop controls. In this way our analysis does not reduce to standard optimal control theory approach and is a separate approach.

Second, by substituting eq. (25) into (26) yields that any interior candidate uninvadable strategy satisfying *s*(*t*, *u*^*^) = 0 (recall 12) must satisfy

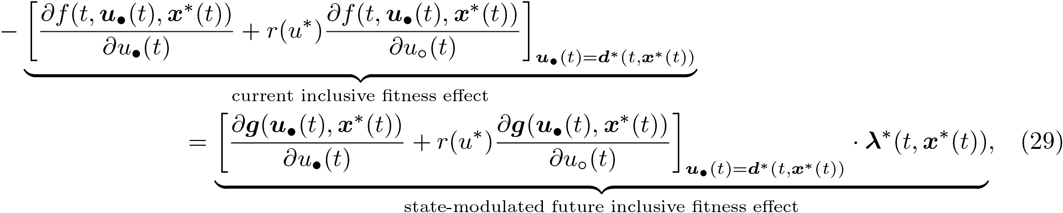

This fundamental balance condition says that *the current inclusive fitness effect* (on the focal individual) is traded-off (hence the negative sign) by the *state-modulated future inclusive fitness effect* resulting from the change in state variables. This trade-off is instrumental in allowing to characterise the candidate uninvadable control *u*^*^(*t*) = *d*^*^(*t*, ***x***^*^(*t*)), which can be typically done in two steps. The first step is to determine *u*^*^(*t*) satisfying (29), while treating the system state ***x***^*^(*t*) and its shadow value **λ**^*^(*t*, ***x***^*^(*t*)) as parameters, yielding the implicit expression

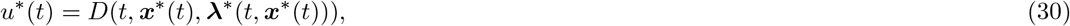

in terms of some function D (that satisfies eq. 29). Essentially, this step is akin to solving a static (onedimensional) first-order condition (and which can in principle also be used whenever *s*(*t*, *u*^*^) = 0, Dockner et al., 2000, p. 97). We will refer to this first-step characterisation as the *static characterisation*, since it allows to characterise the general nature of the solution in terms of ***x***^*^(*t*) and **λ**^*^(*t*, ***x***^*^(*t*)) independently of their explicit values. The second step entails solving for the trajectories of *x*^*^(*t*) and *v*^*^(*t*, ***x***^*^(*t*)) generated by eqs. (27) and (28) under eq. (30) and then taking the gradient of ∇*v*^*^(*t*, ***x***^*^(*t*)) to obtain **λ**^*^(*t*, ***x***^*^(*t*)). Finally, after solving for trajectories *x*^*^(*t*) and **λ**^*^(*t*, ***x***^*^(*t*)) we can explicitly characterise the candidate uninvadable control by substituting these solutions into eq. (30). Solving eq. (28) for *v*^*^(*t*, ***x***^*^(*t*)) is the main technical challenge in finding the candidate uninvadable traits.

It is also often the case in biological models that the Hamiltonian is affine in the control variables so that fitness depends linearly on the evolving traits (e.g. Macevicz and Oster, 1976; Perrin, 1992; Perrin et al., 1993; Irie and Iwasa, 2005; Avila et al., 2019). In such cases, controls do not appear in the selection gradient (26) and hence, one can not directly determine from it the static characterisation (30). These types of controls are known to be *singular arcs* (see Kelley, 1964; Kopp and Moyer, 1965; Goh, 1966 for classic developments and see e.g. Sethi and Thompson, 2006; Bryson and Ho, 1975 for textbook treatments). In order to characterise the candidate uninvadable singular arc, we can take the total time derivative of the selection gradient *s*(*t*, *u*^*^), which (potentially) provides an additional algebraic equation in the variables (*u*^*^, *x*^*^, **λ**^*^) that can contain the control(s) with a non-zero coefficient. In case it does not, another time derivative can be taken until expression for *u*^*^ can be obtained. Hence, for singular arcs, we can obtain the static characterisation (30) by applying

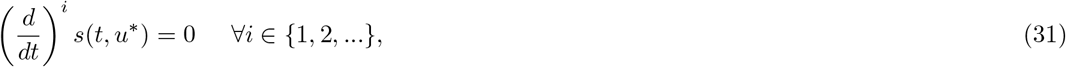

until *u*^*^ can be obtained. Note that for a candidate uninvadable control to be a singular arc, eq. (31) has to hold for a finite interval. If eq. (31) does not hold over a finite interval, then *u*^*^ is known to be a *bang-bang* control (see e.g. Sethi and Thompson, 2006; Bryson and Ho, 1975), meaning that *u*^*^ takes the values only on its boundaries (*u*^*^(*t*) = *u*_max_(*t*) or *u*^*^(*t*) = *u*_min_(*t*), owing to eq. 12).

### 3.2.2 Shadow value dynamics and state feedback in a resident population

From the static (first-step) characterisation of *u*^*^(*t*) = *d*^*^(*t*, ***x***^*^(*t*)) (eq. 30), we observe that the candidate uninvadable trait is at most a function of **λ**^*^(*t*, ***x***^*^(*t*)), but does not directly depend on the reproductive value *v*^*^ itself. Furthermore, taking the partial derivative of eq. (28) with respect to ***x***^*^(*t*) and using the definition of the Hamiltonian (25) and re-arranging (see sections B.1.2 and B.3 in Appendix) yields

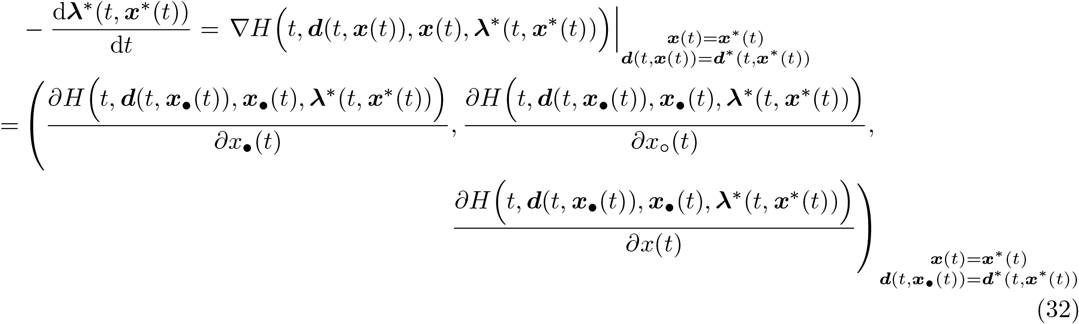

with f.c. **λ**^*^(*t*_f_, ***x***^*^(*t*_f_)) = ∇Φ(***x***^*^(*t*_f_)) and where ***d***(*t*, ***x***_∂_(*t*)) = (*d*(*t*, ***x***_o_(*t*)), *d*(*t*, ***x***_o_(*t*), *d*(*t*, ***x***(*t*)). The dynamics of the shadow value, given by eq. (32), may at first glance appear to be an ODE (and therefore easier to solve than eq. 28, which is clearly a PDE for the reproductive value *v*^*^). This may lead one to hope that it is possible to circumvent from explicitly determining *v*^*^, by simply solving eq. (32) to directly obtain **λ**^*^. But this hope is crushed by the trait dependence on state, which entails that eq. (32) depends on the derivatives of the elements of ***d***(*t*, ***x***_•_(*t*)) with respect to state, which in turn depends on higher-order derivatives of *v*(*t*, ***x***^*^(*t*); *u*^*^). This can be seen by using eq. (30), whereby

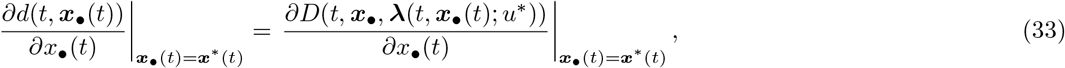

which unveils that eq. (32) is actually a PDE. This means that in general it is not possible to determine the candidate uninvadable trait from using eq. (32), which has been repeatedly stressed in optimal control theory (e.g. Starr and Ho, 1969*a*, *b*; Başar, 1977). However, the analysis of the components of eq. (32) has less been stressed, but turns out to be informative in highlighting the main similarities and differences between selection on closed-loop and open-loop controls.

Lets now decompose eq. (32) for the component 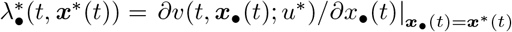 (similar results hold for the other shadow values (see Appendix B.3) and write

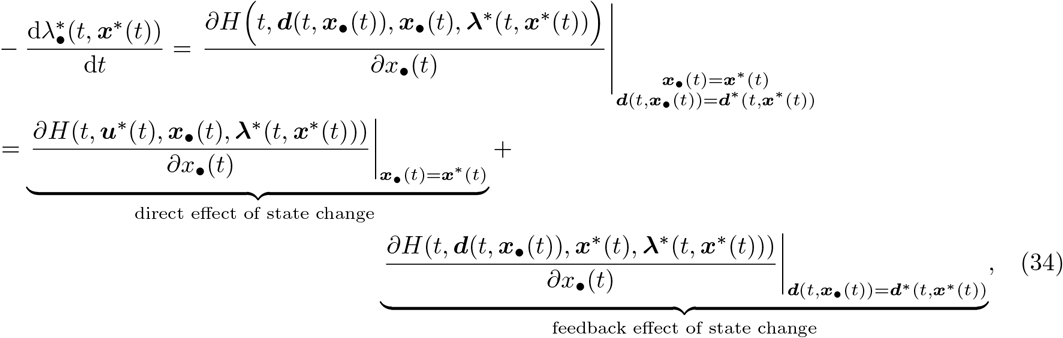

with f.c. 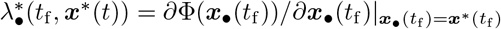. This says that the rate of change of the shadow value is given by a direct effect of state change on the Hamiltonian (current fitness effect and state-modulated fitness effect) and a feedback effect on the Hamiltonian, which arises since closed-loop traits react to changes in the state. Using the expression for the Hamiltonian in eq. (25), the expressions for direct and indirect effects in eq. (23)–(24), and noting that *∂d*(*t*, ***x***(*t*))/*∂x*_•_(*t*) = 0, the trait feedback effect can be further expanded as

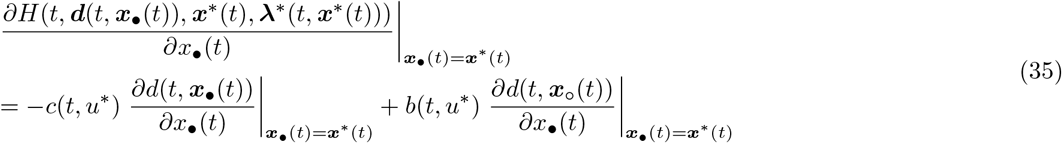

where the derivatives *∂d*(*t*, ***x***_•_(*t*))/*∂x*_•_(*t*) and *∂d*(*t*, ***x***_o_(*t*))/*∂x*_•_(*t*) give the trait sensitivities of the focal individual and its average neighbour, respectively, at time t to changes in focal’s state variable *x*_•_(*t*). Hence, the feedback effect of state change is equal to the trait sensitivities of all individuals in the group weighted by their effects on the focal’s fitness (the latter are effectively the direct and indirect fitness effects).

We now make three observations about eqs. (34)–(35). First, trait sensitivities result in inter-temporal feedbacks in trait expressions. We can see this by first observing that current trait expression affects changes in state variables (by way of the second line of eq. 29) which affect future fitness (measured by the shadow value). In turn, the dynamics of shadow value takes into account that closed-loop traits respond to changes in state variables (by way of the second line of eq. 35). That is, the shadow value takes into account the effects of current trait expression on future trait expression. Hence, under closed-loop control, the current trait expression of one individual is linked to future trait expression of itself and other individuals in its group. Second, the sign of the feedback effect of state change (sign of eq. 35) determines the direction of the effect of trait sensitivities on the shadow values. This means that the sign of the feedback effect balances the trade-off between current and future (state-modulated) fitness effects (by way of eq. 29). For positive feedback effect, trait sensitivity increases future (inclusive) fitness gains (second line of eq. 29), while for negative feedback effect, trait sensitivity decreases future (inclusive) fitness gains. Third, the shadow value dynamics given by eqs. (34)–(35) is different from that in classical results from dynamic game theory (first developed by Starr and Ho, 1969a, b), where the feedback effect through the focal’s own trait variation does not appear due to the absence of interactions between relatives, whereby –*c*(*t*, *u*^*^) = 0 at *s*(*t*, *u*^*^) = 0. By contrast, in our model with interactions between relatives one has –*c*(*t*, *u*^*^) + *r*(*u*^*^)*b*(*t*, *u*^*^) = 0 at *s*(*t*, *u*^*^) = 0. Thus, we recover the classical result for the feedback effect from dynamic game theory when *r*(*u*^*^) = 0.

We now consider three scenarios (which are relevant for biology) under which the feedback term (given by eq. 35) that describes the dynamics of 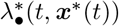 vanishes (similar arguments also hold for the feedback term 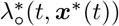 and recall that we do not need to consider the dynamics λ^*^(*t*, ***x***^*^(*t*)) here, because λ^*^(*t*, ***x***^*^(*t*)) does not affect the selection gradient). That is, we consider scenarios for which eq. (32) is a system of ODE’s (for components 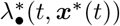 and 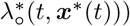 and therefore solving a PDE (28) for *v*^*^(*t*, ***x***^*^(*t*)) is not necessary to determine the candidate uninvadable trait. These three scenarios are as follows.

i. Open-loop *u*(*t*) = *d*^*^(*t*) controls. Because the traits do not depend on the state variables, *∂d*_•_(*t*)/*∂x*_•_(*t*) = 0 and *∂d*_o_(*t*)/*∂x*_•_(*t*) = 0, which implies that eq. (35) vanishes.
ii. No social interactions in the group, meaning that fitness components of individuals do not depend on traits and states of other individuals in the group, i.e. the fitness components *f*, Φ and *g* of the focal individual do not depend on *u*_o_ and *x*_o_, hence *b*(*t*, *u*^*^) = 0. It then further follows from eq. (26) that in order for *s*(*t*, *u*^*^) = 0 to be satisfied, we need *c*(*t*, *u*^*^) = 0. It then follows directly that eq. (35) vanishes.
iii. In a population of clonal groups (*r*(*u*_*_) = 1) that share a common state variable (*x*_•_(*t*) = *x*_o_(*t*), e.g. common resource in the group). Two observations can be made for this scenario. First, from eq. (26) it follows that –*c*(*t*, *u*^*^) + *b*(*t*, *u*^*^) = 0 for clones at *s*(*t*, *u*^*^) = 0. Second, since *x*_•_(*t*) = *x*_o_(*t*), then e.g. *∂d*(*t*, ***x***_•_(*t*))/*∂x*_•_(*t*) = *∂d*(*t*, ***x***_o_(*t*))/*∂x*_•_(*t*). Combining these two observations directly leads to conclude that the feedback term eq. (35) vanishes.

There are a two implications that follow for these three cases. First, open-loop and closed-loop evolutionary equilibria are in general different, since the state-feedback effect causes inter-temporal feedbacks between trait expressions of locally interacting individuals under closed-loop controls, which are not possible under open-loop controls. However, if individuals do not locally interact or if they interact in clonal groups, then closed-loop and open-loop representation of controls produces the same candidate uninvadable trait and state trajectories. Second, since the feedback effect (eq. 35) vanishes for open-loop controls, the sign of the feedback effect is crucial in comparing closed-loop and open-loop controls. Most importantly, the sign of the feedback effect allows to compare the balance of the trade-off between current versus future (inclusive) fitness effects between open-loop and closed-loop controls (by way of eq. 29). If the feedback effect is positive, then the (inclusive) fitness gain from future (second line of eq. 29) is higher under closed-loop control than under open-loop control. If the feedback effect is negative, then the (inclusive) fitness gain from future is lower under closed-loop control than under open-loop control.

### 3.3 First-order conditions for open-loop controls

We now focus specifically on open-loop controls by pointing out the simplifications that arise when the decision rule depends only on time:

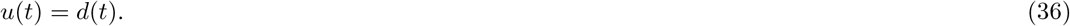

As we showed in the previous section, for open-loop controls the state-feedback term in eq. (35) for the dynamics of the shadow value 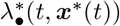 vanishes since *∂d*_•_(*t*)/*∂x*_•_(*t*) = 0 and *∂d*_o_(*t*)/*∂x*_•_(*t*) = 0 (similarly it vanishes also for 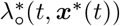 and λ^*^(*t*, ***x***^*^(*t*))), which implies that eq. (32) is a system of ODE’s). Hence, we can characterise the necessary condition for an open-loop control paths as follows.

### Open-loop control result

*Let* 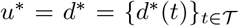 *be the candidate uninvadable open-loop control path with associated state path* 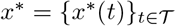 *and shadow value* **λ**^*^(*t*) = **λ**^*^(*t*, ***x***^*^(*t*)). *The candidate uninvadable control path u*^*^ = *d*^*^ *has to necessarily satisfy* *eq.* (12), *where the point-wise selection coefficient s*(*t*, *u*^*^) *on a control component u*^*^(*t*) = *d*^*^(*t*) *can be written for all* 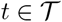

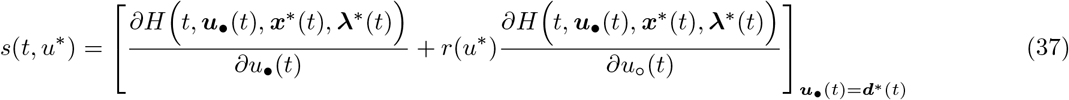

*where* ***d***^*^(*t*) = (*d*^*^(*t*), *d*^*^(*t*), *d*^*^(*t*)), *the state variable satisfies*

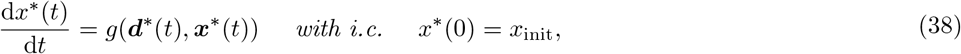

*and the shadow values satisfy*

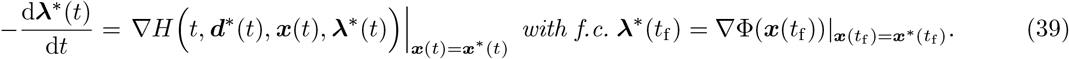

This result is Pontryagin’s weak principle for interactions between relatives (since only small mutant deviations are considered, Speyer and Jacobson, 2010, p. 74) and only requires consideration of the shadow value. It has been derived previously Day and Taylor (1997, 2000), for a slightly less general model, where individuals locally play the field (fitness only depends on the traits of individuals in the focal group) or interact in a pairwise way (see also Day and Taylor, 1998; Wild, 2011 for related work). Yet this result covers both group-structured and panmictic populations and thus covers the first-order condition result of Metz et al. (2016) as well as those of classical life-history models (e.g., Perrin and Sibly, 1993 for a review). We here re-derived this result as a corollary of the closed-loop result of the previous section when the feedback-terms describing the dynamics of the shadow value **λ**(*t*) vanish (i.e, eq. 35 vanishes). Hence, we closed the loop between the selection gradient on function-valued traits, invasion implies substitution, Hamilton’s rule, dynamic programming, and Pontryagin’s (weak) maximum principle.

### 3.4 Special case controls

We now work out further simplifications that arises in the characterisation of the first-order condition when more specific biological assumptions are made.

#### 3.4.1 Stationary controls

For instance, under several biological situations, the individual fitness function (eq. 13) can be expressed in the simpler form

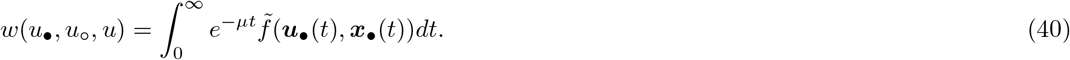

Here, the time horizon is large (*t*_f_ → ∞), there is no scrap value (Φ(***x***_•_(*t*_f_)) = 0), μ is a constant mortality (or discount) rate so that *e*^−μ*t*^ can be interpreted as the probability of survival until time *t*, and 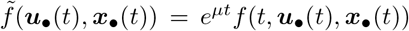 is the current rate of fitness increase at time *t*, which is assumed to not explicitly dependent on the time variable *t* (in game theory, fitness functions like eq. 40 cover the so-called infinite-horizon autonomous differential games, Dockner et al., 2000; Weber, 2011).

The relevant feature of this setting is that, conditional on reaching a certain time (or age), the remaining fitness is the same as the original fitness; future fitness is thus time invariant. This can be seen more explicitly by considering the reproductive value. Indeed, conditional on reaching time *t*, which occurs with probability *e*^−μ*t*^, the conditional reproductive value starting at *t*–*the current-value reproductive value*–is given by

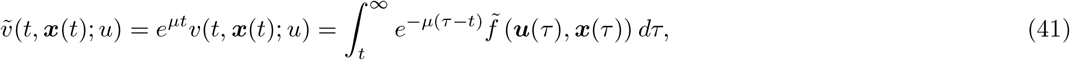

and this is Fisher’s reproductive value under the present assumptions (e.g., Goodman, 1982; Charlesworth, 1994). It takes exactly the same functional form starting at any time 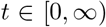 (in game-theory terms it is said that “fundamentals of the game do not change over time” Dockner et al., 2000, p. 97). Owing to this time invariance, trait expression could be taken to be time invariant as well and we now let the control take the following form

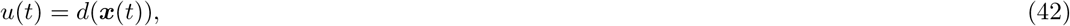

which we call a (closed-loop) *stationary* control (called a stationary feedback control or a stationary Markov strategy in game theory, e.g., Dockner et al., 2000; Weber, 2011). On substituting the stationary control (42) into eq. (41), we see that the current-value reproductive value is independent of time and thus stationary

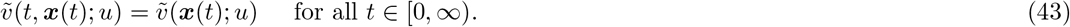

This means that given some fixed initial value ***x***(*t*) = ***x***_init_, the current-value reproductive value is the same regardless of the time the process is started. So any variation in the initial value has a timeindependent effect on 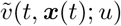, which, more formally owing to eq. (43) satisfies

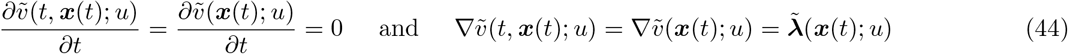

where 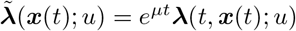 is the current-value shadow value that depends on time only indirectly through ***x***(*t*).

This feature of stationary controls allows to markedly simplify their first-order characterisation (as shown by many examples of the game theory literature e.g., Dockner et al., 2000; Weber, 2011, p. 97). In order to carry out this characterisation explicitly it is useful to use the notion of current-value Hamiltonian (Weber, 2011, p. 111), which in our setting is defined as

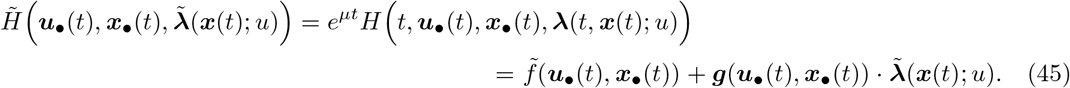

Applying eqs. (41)–(45) to eqs. (26)–(28) establishes the necessary condition for uninvadability of stationary controls as follows.

##### Stationary (closed-loop) control result

*Let* 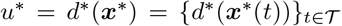 *be a candidate uninvadable stationary (closed-loop) control path with associated state path* 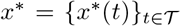 *and stationary current-value shadow value* 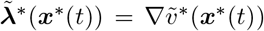 *with current-value reproductive value* 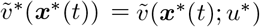. *The candidate uninvadable control path d*\*(***x***\*) *has to necessarily satisfy* *eq*. (12), *where the point-wise selection coefficient s*(*t,u*\*) *on control component u*\*(*t*) = *d*\*(*x*\*(*t*)) *can be written for all* 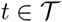 *as*

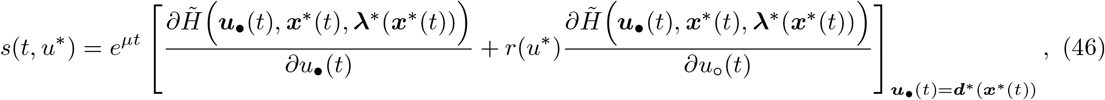

*where **d***^*^(***x***^*^(*t*)) = (*d*^*^(***x***^*^(*t*)), *d*^*^(***x***^*^(*t*)), *d*^*^(***x***^*^(*t*))), *the state variable satisfies*

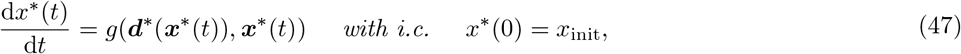

*and the current-value reproductive value satisfies*

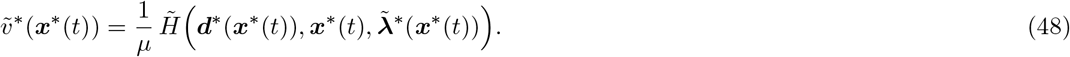

Here, eq. (48) follows from taking the (partial) time derivative of (first two equalities of) eq. (41) and using eq. (44) to obtain 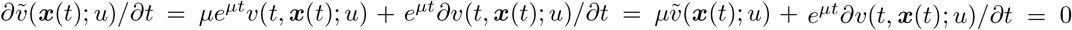 and then using eqs. (28) and (45). Since, 1/*μ* is the average length of an interaction (or lifespan), eq. (48) says that total expected fitness 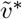 is equal to the expected fitness gain 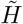 from current phenotypic expression (a fitness flow) multiplied by the average length of time that fitness gain can accrue (and this holds for each state). By contrast to eq. (28), eq. (48) does not involve any partial derivatives with respect to time. Furthermore, if ***x***\*(*t*) is one-dimensional then 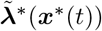 is also one-dimensional, and in this case eq. (48) is an ordinary differential equation. Hence, the analysis of stationary controls is generally more tractable.

#### 3.4.2 Constant controls

We finally turn to a type of control that that is not analysed in the optimal control theory literature, yet is relevant to evolutionary biology. This is the case of *constant control* 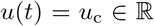 for all 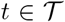, where the subscript c emphasises that the control is independent of time. While constant controls are essentially scalar traits, these traits may nevertheless affect the dynamics of state variables that in turn affect fitness, and this makes the analysis of a necessary first-condition for uninvadability time-dependent. We finally provide this characterisation as a corollary of the “Open-loop control result” result as follows.

##### Constant control result

*Let* 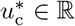 *be a candidate uninvadable constant control (a scalar trait) with associated state path* 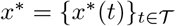 *and shadow value* **λ**^*^(*t*). *The candidate uninvadable control u*^*^ *has to satisfy* *eq*. (10) *with the (scalar) selection coefficient*

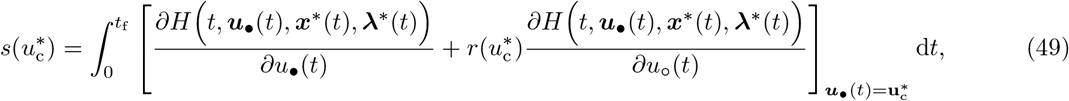

*where* 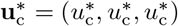, *the state variable satisfies*

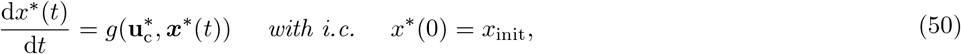

*and the shadow values satisfy*

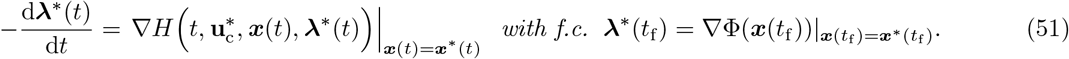

The key difference between this result and the “Open-loop control result” is eq. (49), which says that the selection coefficient depends on the derivatives of the Hamiltonian integrated over the interaction period 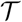. This follows directly from the fact that the control is a scalar and a mutant schedule is deviated in the same direction at all time points (hence there are no differences in point-wise deviations). Rather surprisingly, this or closely related results do not seem to have appeared previously in the optimal control nor evolutionary biology literature, even for the special cases of no interactions between individuals. However, this result may be useful for example in connecting our approach with various previous models (see Discussion and Box 1 for a concrete example).

## 4 Examples

### 4.1 Common pool resource production

#### 4.1.1 Biological scenario

We here present an application of our results to the production of common pool resource that returns fitness benefits to all group members but is produced at a personal fitness cost. The evolving trait we consider is the schedule 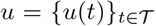 of effort invested into resource production during an interaction period 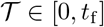 and we will hereinafter refer to the control *u*(*t*) as the *production effort* at time *t*. Let *u*_•_(*t*), *u*_o_(*t*), *u*(*t*) denote the production efforts at time *t* of the focal individual, average neighbour in the focal group, and average individual (from other groups) in the population, respectively. Let *x_c_*(*t*) and *x*(*t*) be the *resource level* at time *t* (the total amount of resources produced) in the focal group and the average group in the population, respectively. Note that here we have a common state variable *x*_c_(*t*) = *x*_•_(*t*) = *x*_o_(*t*) between individuals in the same group and individuals interact through the state only locally (resources produced in other groups does not directly affect the fitness of individuals in the focal group).

We study the evolution of the production effort under two different scenarios: (i) individuals can adjust their production effort according to the (local) resource level *x_c_*(*t*) (closed-loop control) and (ii) individuals are committed to a fixed schedule of production effort (open-loop control). One difficulty in analysing the evolution of such traits is that limited genetic mixing generates relatedness between group members but also competition between them, which leads to kin competition (e.g., Taylor, 1988; Frank, 1998; Rousset, 2004; Van Cleve, 2015). Since we want to highlights the key effects of the evolution of open-loop and closed-loop controls in the context of interactions between relatives in a simple way, we want to avoid the complexities brought by kin competition and thus assume implicitly a life cycle that entails no kin competition and that relatedness is independent of the control *r*(*u*) = *r*.

In particular, we assume that individual fitness takes the form

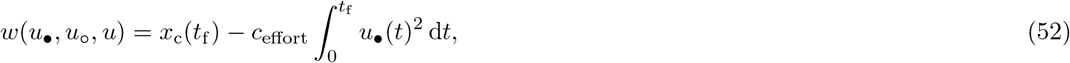

which depends on the resource level *x*_c_(*t*_f_) in the group at the end of interaction period *t*_f_ and on the accumulated (personal) cost of producing the common resource for the group (the second term in eq. 52). Here *c*_effort_ is a parameter scaling the personal cost. The resource level *x*_c_(*t*_f_) and *c*_effort_ are measured in units of the number of offspring to the focal individual and scaled such that they inherently take into account the proportional effect of density-dependent competition (proportional scaling of fitness does not affect the direction of selection). We assume that *x*_c_(*t*) depends on the total amount of production effort that individuals in the focal’s group invest into producing it and that the return from this effort decreases exponentially with the current level of resource

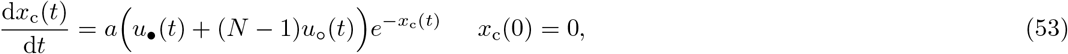

where the parameter *a* > 0 is the efficiency of producing the common resource.

We now make two observation about this model. First, neglecting the effects of kin competition in eq. (52) does not lead to any loss of generality in our forthcoming analysis, since taking kin competition into account would only affect the final results by re-scaling the value of relatedness (e.g., Van Cleve, 2015; Mullon et al., 2016). Second, the mathematical properties and thus the structure of the game embodied in eqs. (52)–(53) are equivalent to those of the parental investment game model of Ewald et al. (2007) from which we took inspiration. However, biologically eqs. (52)–(53) have a different interpretation, since our model considers interactions between relatives (Ewald et al., 2007 considers pair-wise interactions of non-relatives) and we will compares open-loop and closed-loop controls (Ewald et al., 2007 compares static traits with closed-loop traits). The forthcoming analysis will also conceptually depart from that of Ewald et al. (2007), because it will be based on the (neutral) reproductive value function (28), while their analysis is based on solving the Hamilton-Jacobi-Bellman equation for the optimal value function (see Ewald et al., 2007, p. 1456) with which the present n— player scenario cannot be obtained as a trivial extension of the approach used in (Ewald et al., 2007).

#### 4.1.2 Static characterisation of the production effort

From eq. (52), the reproductive value (eq. 19) for this model is

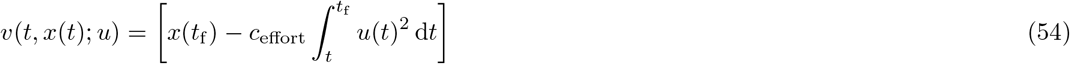

We denote the corresponding shadow value with λ_c_(*t,x*; *u*) = *∂v*(*t, x*_c_(*t*);*u*)/*∂x*_c_(*t*)|_*x*_c_(*t*)=*x*(*t*)_. Eq. (52) also entails that the fitness components *f* and Φ (as defined in eq. 13) take the form

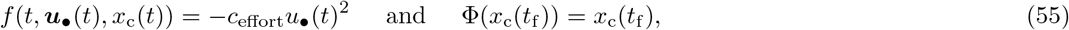

while the rate of change of the state variable *x*_c_(*t*) is given by

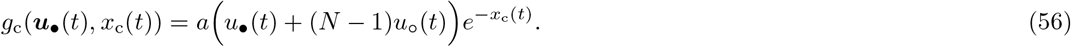

On substituting eqs. (55)–(56) into the Hamiltonian (25) produces

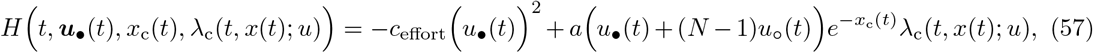

which gives the direct fitness effect

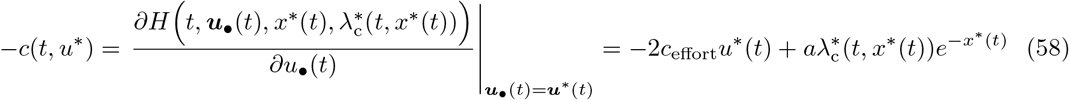

and the indirect fitness effect

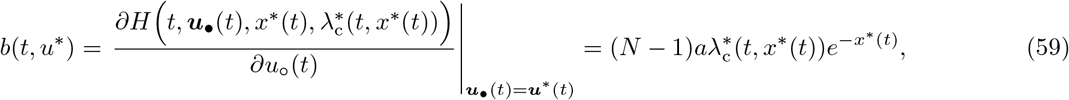

where 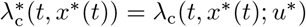. Hence, the balance condition (29) for this model reads

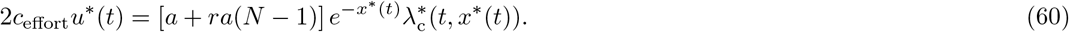

This says that the net effect on accumulated personal cost due to spending effort to produce a unit resource must balance out the inclusive fitness benefit associated with that unit resource. Solving eq. (60) for *u*^*^(*t*) yields

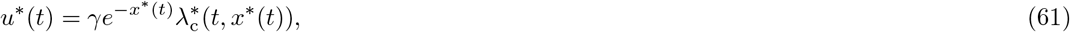

where

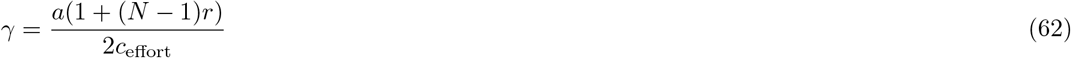

scales the benefit to cost ratio of producing the resource, note also that *γ* > 0. Eq. (61) says that (candidate uninvadable) production effort *u*^*^(*t*) decreases exponentially with the resource level *x*^*^(*t*), increases linearly with the shadow value and relatedness, and is not directly dependent on time. This general nature of the solution applies to both open-loop and closed-loop controls and is depicted in Fig. 2.

**Figure 2:**
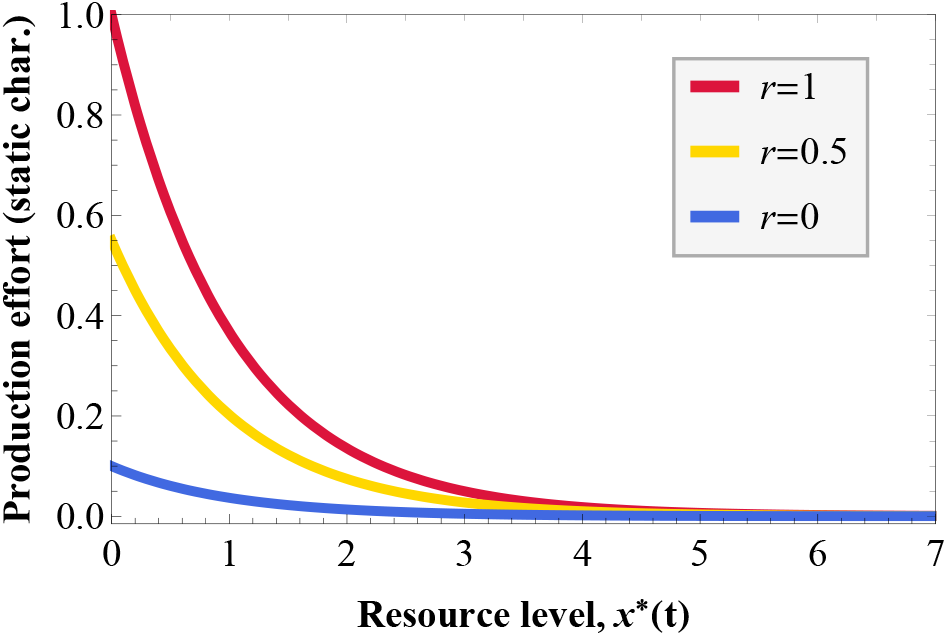
Static characterisation 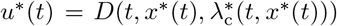 (eq. 61) of the candidate uninvadable production effort as a function of resource *x*\*(*t*) for fixed values of 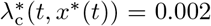 and for different values of relatedness between individuals in the group. Parameter values: *N* =10, *a* =1, *c*_effort_ = 0.01. Note that characterisation holds for both open-loop and closed-loop controls.

#### 4.1.3 Closed-loop production effort

We now turn to analyse *u*\*(*t*) explicitly as a function of time when the control is the closed-loop *u*\*(*t*) = *d*^*^(*t, x*^*^(*t*)), which requires to evaluate *x*^*^(*t*) and *v*^*^(*t, x*^*^(*t*)) = *v*(*t, x*^*^(*t*); *u*^*^). To that end, we evaluate the dynamic eq. (53) for *x*_c_(*t*) along ***u***_•_ = ***u***^*^ and *x*_c_ = *x*^*^ and substituting the expression for *u*^*^(*t*) from eq. (61), whereby

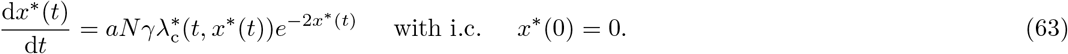

Substituting eq (57) and eq. (61) into eq. (28) and simplifying yields

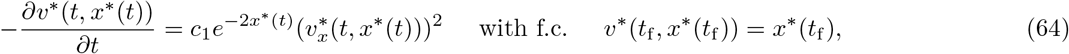

where

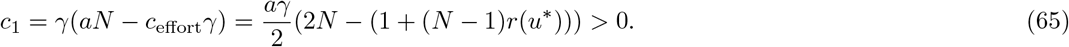

Using the method of characteristics, Ewald et al. (2007) showed that the partial differential equation (64) has the following solution

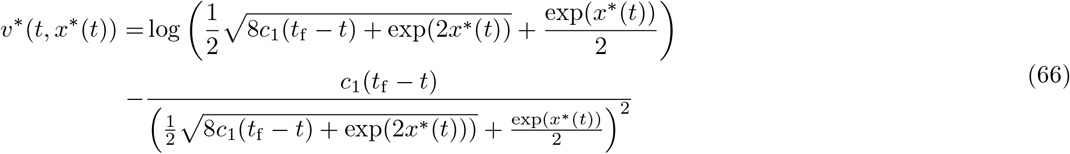

(our eq. 64 corresponds to eq. (21) of Ewald et al., 2007 where *c*_1_ = 3/2*k* and their solution is presented on page 1459 of their paper, where *c*_1_ = *c* = 3/2*k*). Taking the derivative with respect to *x*\*(*t*) and upon simplifying yields the expression for the shadow value

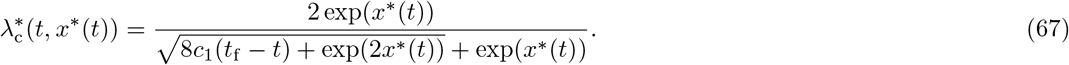

Substituting this into the static characterisation eq. (61) shows that

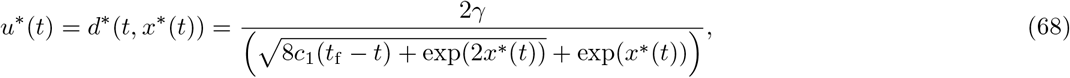

where the state variables is the solution of

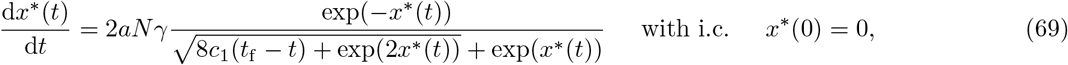

which was obtained by substituting eq. (67) into eq. (63) and for which we were not able to find a closed form expression.

#### 4.1.4 Open-loop production effort

We turn to derive the candidate uninvadable open-loop trait *u*\*(*t*) = *d*\*(*t*). Substituting eq. (57) and eq. (61) into eq. (39), we arrive, by using eq. (63), at the following two-point boundary value system

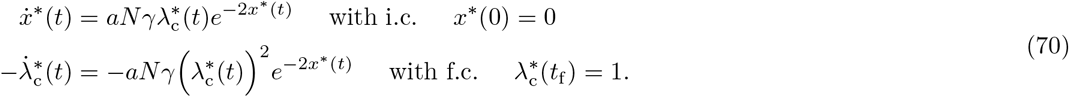

This system has one real-valued solution

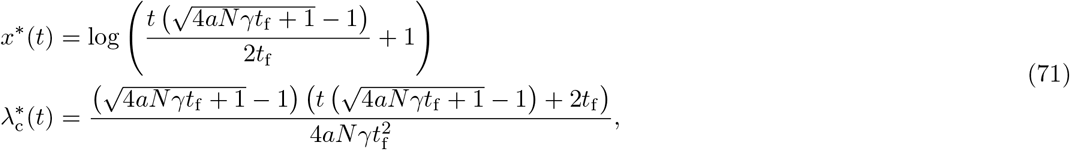

and substituting this solution back into eq. (61) produces

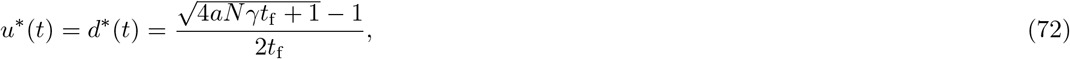

which turns out to be constant in time.

#### 4.1.5 Comparison between closed-loop and open-loop production efforts

This example illustrates our generic result that in a population of clonal groups (*r* = 1) closed-loop and open-loop equilibria coincide (Figure 3). In a population of non-clonal groups (*r* < 1) the production effort *u*^*^(*t*) and the resulting amount of resource *x*^*^(*t*) tend to be lower under the closed-loop equilibrium (hereinafter, we simply write “equilibrium”) than under the open-loop equilibrium (Figure 3). Overall, the production effort monotonically increases over time for the closed-loop control and stays constant under the open-loop control (Figure 3).

**Figure 3:**
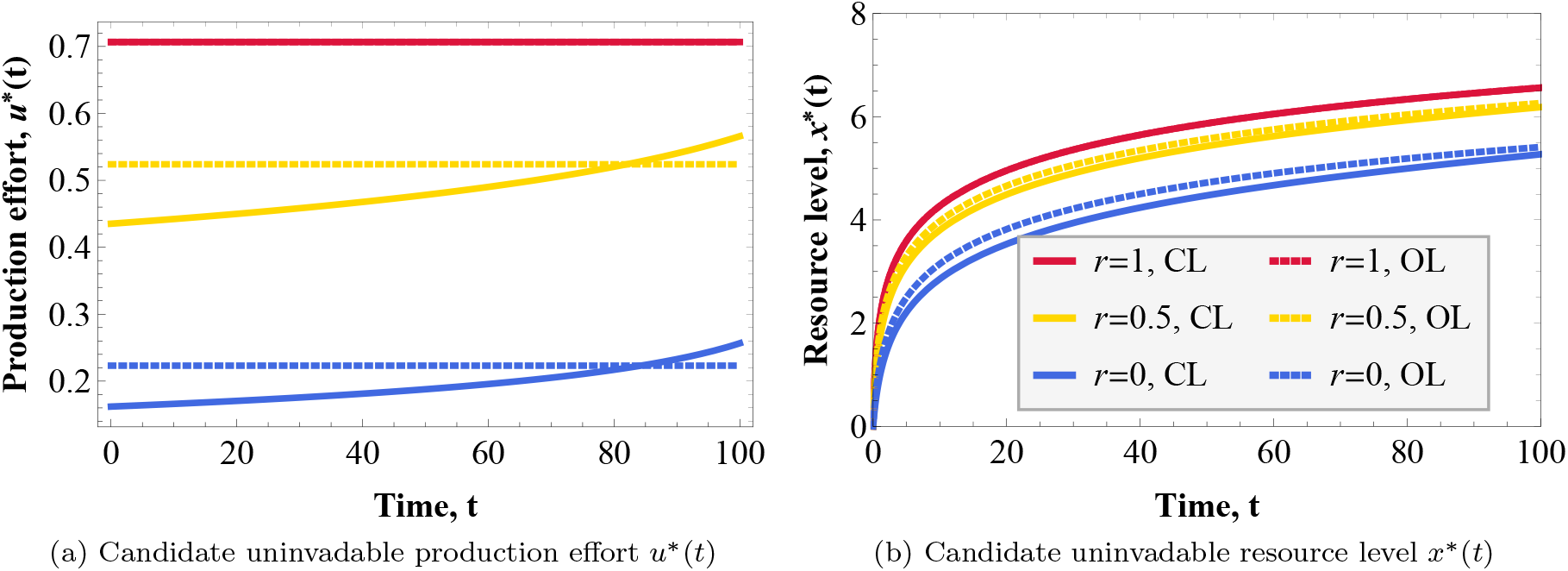
Candidate uninvadable production effort (panel a) and resource level (panel b) for closed-loop (CL) (solid lines) and open-loop (OL) traits (dashed lines) for different values of average relatedness *r*. Parameter values: *N* = 10, *a* =1, *c*_effort_ = 0.01, *T* = 100. Note that if individuals in the group are clones (*r* = 1), the closed-loop and open-loop traits coincide.

The difference between closed-loop and open-loop (non-clonal) equilibria arises from the difference in the shadow value dynamics (recall section 3.2.2). We find that the shadow value is lower under closed-loop control than under open-loop control when *r* < 1 (Fig. 4). This is so because of the feedback effect of state change (eq. 35), which for our example is

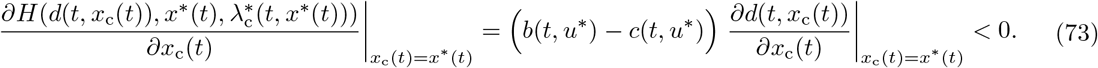

**Figure 4:**
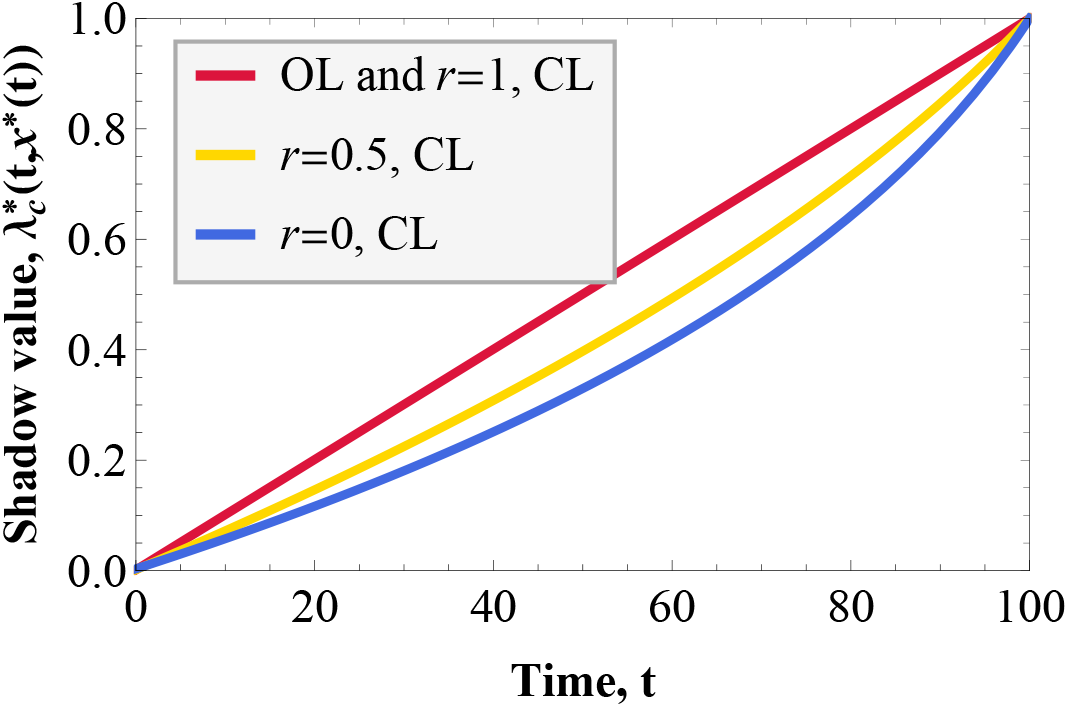
Shadow value 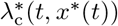 over time t for closed-loop (CL) and open-loop (OL) control and for different values of relatedness between individuals in the group. Parameter values: *N* =10, *a* =1, *c*_effort_ = 0.01, *T* = 100. Note that characterisation holds for both open-loop and closed-loop characterisation of the trait expression rule.

Since this is negative, the shadow value declines faster backwards in time than under the closed-loop equilibrium. In order to understand why the feedback effect is negative, we need to consider the signs of *b*(*t, u*\*) – *c*(*t,u*\*) and the trait sensitivity *∂d*(*t,x*_c_(*t*))/*∂x*_c_(*t*) (which is here the same for all group members). The term *b*(*t,u*\*) – *c*(*t,u*\*) is positive when groups are non-clonal and zero when they are clonal (Fig. 5, panel a). This means that if everyone in the group produces more of the resource, then the focal’s fitness increases under the non-clonal equilibrium and is unaffected under the clonal equilibrium. The trait sensitivity is always negative (Fig. 5, panel b). Hence, individuals will reduce their production effort in response to an increase in the resource level and the magnitude of this effect is larger for higher values of relatedness.

**Figure 5:**
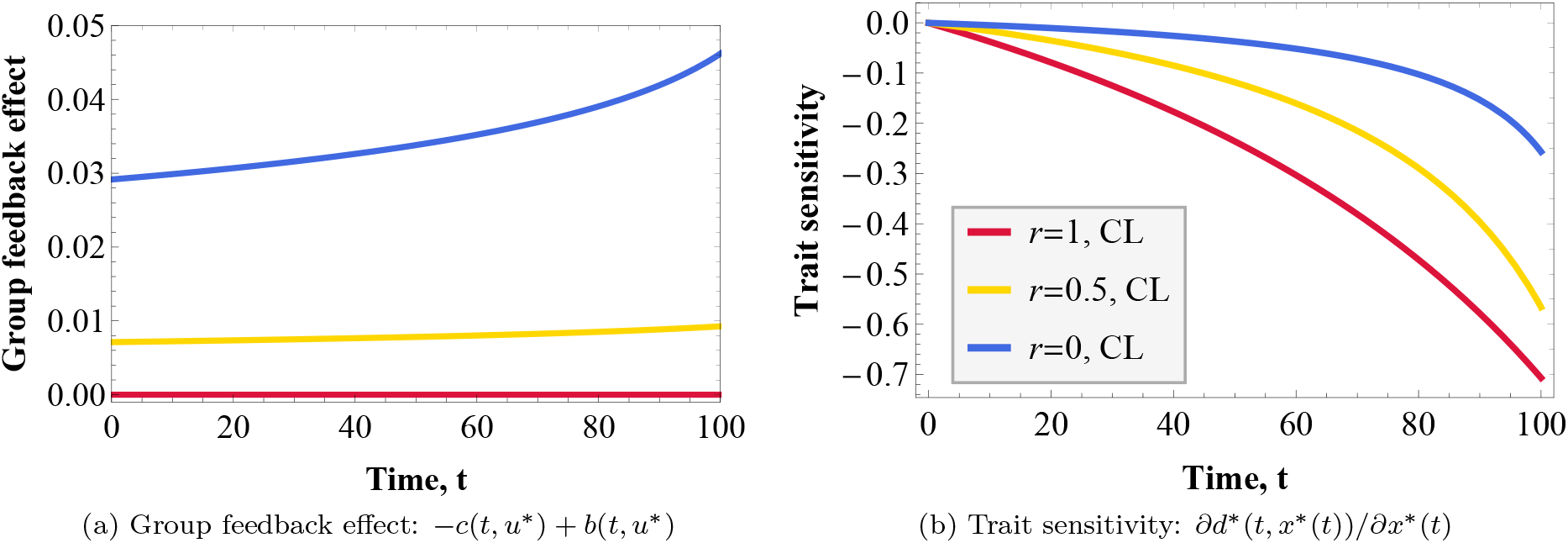
Group feedback effect on focal’s fitness (panel a) and trait sensitivity (panel b) for closed-loop (CL) control for different values of average relatedness *r* in the group. Parameter values: *N* =10, *a* =1, Ceffort = 0.01, *T* = 100.

In conclusion, investment effort is lower for closed-loop traits than for open-loop traits in a population of non-clonal groups (*r* < 1), because closed-loop trait expression takes into account that other individuals will reduce their production effort in response to the focal individual increasing its production effort. In a population of clonal groups where the focal individual increases its trait expression, the response from other individuals will not affect the fitness returns to the focal. For open-loop controls, the response from other individuals can never affect the fitness returns, because open-loop control trajectories are pre-determined at birth (full commitment to control trajectory) and trait expression can not be adjusted in response to changes in the resource level. For clonal groups, trait sensitivity to changes in the resource level does not alter individual behaviour, because everyone’s interests in the group are aligned.

### 4.2 Common pool resource extraction

#### 4.2.1 Biological scenario

We finally present an application for a stationary (closed-loop) control, which involves the same demographic assumption as the previous example but with fitness given by eq. (40). And we now let *u*_•_(*t*), *u*_o_(*t*), *u*(*t*) denote the extraction rates of a given resource with abundance *x*_c_(*t*) at time *t* in the focal group, which is again taken as a common state variable among group members (*x*_c_(*t*) = *x*_•_(*t*) = *x*_o_(*t*)). This resource is assumed to follow the dynamics

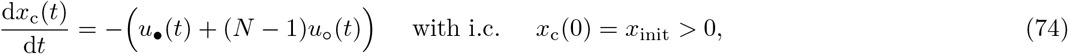

and is thus depleted at a rate given by the sum of the extraction rates within the focal’s group. We assume that the resources extracted by the focal individual translates into current-value fitness according to

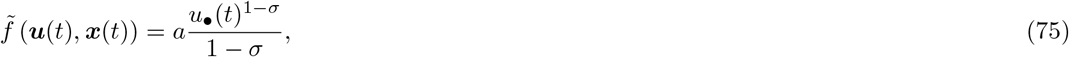

where the parameter *a* > 0 is the efficiency of turning extracted resources into producing offspring and *σ* ∈ (0,1) is a parameter shaping the concavity of the production function, which thus exhibits diminishing returns (we used eq. 75 instead of the perhaps biologically more realistic Holling type 2 functional response for ease of calculations). The model defined by eqs. (74)–(75) recasts into the homogeneous island model the standard baseline resource extraction game of environmental economics (Dockner et al., 2000, chapter 12.1, Weber, 2011, chapter 4.3).

#### 4.2.2 Static characterisation of the extraction rate

On substituting eqs. (74)–(75) into the current-value Hamiltonian (45) and considering only a current-value shadow value 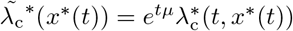 (since we have common state variable) produces

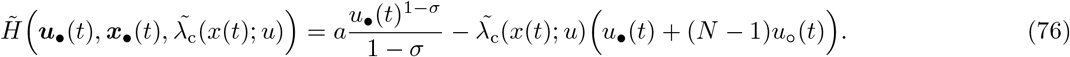

This yields the direct fitness effect

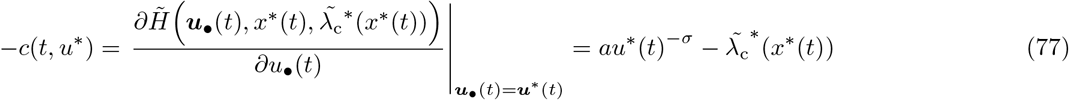

and the indirect fitness effect

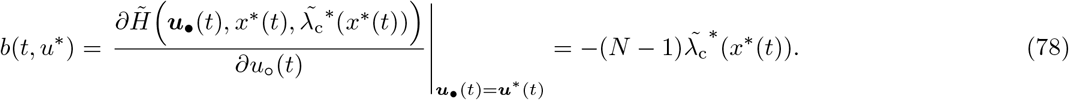

For this model, the balance condition reads

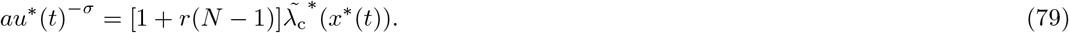

This says that the net present personal benefit of a unit extracted resource must balance out the future inclusive fitness cost resulting from depleting that unit resource, and yields the equilibrium extraction rate

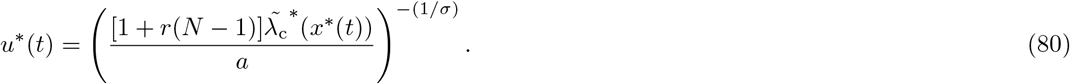

#### 4.2.3 Stationary (closed-loop) extraction rate

Let us now analyse *u*^*^(*t*) explicitly as a function of time when the control is stationary *u*^*^ (*t*) = *d*^*^ (*x*^*^ (*t*)). Substituting eq. (76) and eq. (80) into eq. (48), simplifying and dropping the time index on the state variables yields

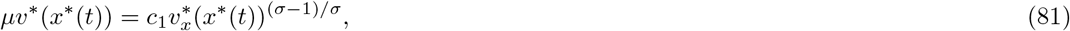

which is a differential equation for *v*\*(*x*\*) where 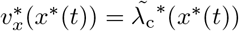 and

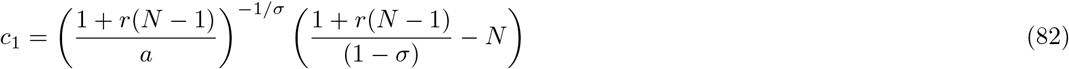

is a constant that we assume to be positive (which thus requires that 1 + *r*(N – 1) > *N*(1 – *σ*)). Following Dockner et al. (2000, p. 325) and arguments therein, the solution to eq. (81) under the given constraints is

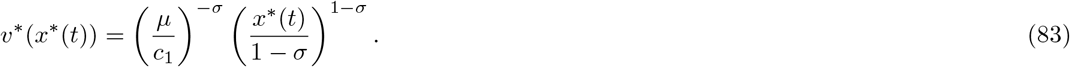

Owing to the fact that 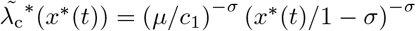, the stationary control (80) written explicitly as a function of time becomes

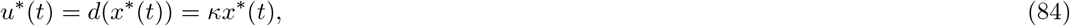

where

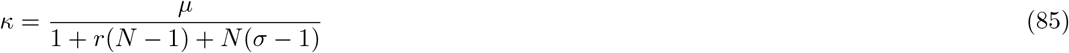

can be interpreted as the equilibrium extraction rate of a unit resource by an individual, since *x*^*^(*t*) such units are available at *t*. Substituting eq. (84) into eq. (74) and solving the latter gives the candidate uninvadable control and state path explicitly as

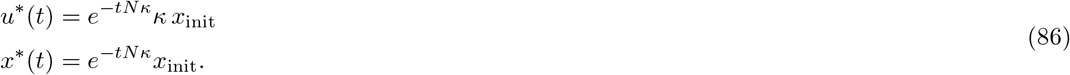

When *r* = 0, we thus recover the standard result for the stationary control established in the game theory literature [the symmetric stationary Markov Nash equilibrium (Dockner et al., 2000, eq. 12.38)], while when *r* = 1 we recover the game theory cooperative solution (Dockner et al., 2000, eq. 12.7).

#### 4.2.4 Open-loop extraction rate

We now work out an open-loop control for this model and to this end it is useful to express the static characterisation (80) in terms of the original shadow value 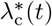 (not the current value shadow value 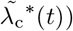, which, on recalling that 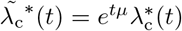, yields

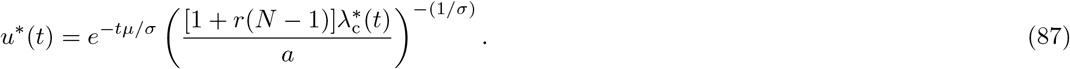

Note that 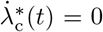 in eq. (39) because the partial derivative of the current-value Hamiltonian 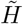 with respect to the second argument is zero (by way of eq. 76) and recall also that 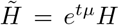 (hence the partial derivative of Hamiltonian with respect to its third argument is also zero). From this it follows that 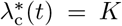 for some constant *K* > 0 (positive since otherwise no resources will be extracted). Inserting this into eq. (87) and then substituting the resulting expression for *u*^*^(*t*) into the state dynamic (74) gives

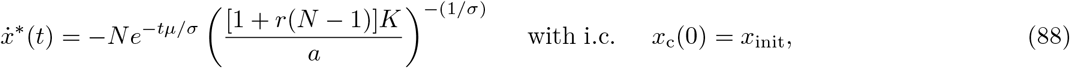

whose solution is

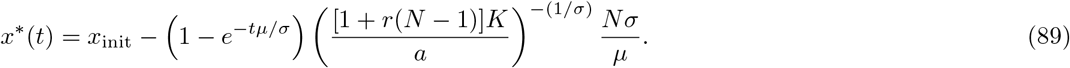

We now seek an open-loop solution *u*^*^(*t*) that satisfies

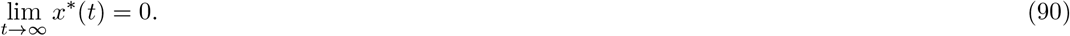

That is, we assume that all the resources will be depleted asymptotically. In game-theory terms, this corresponds to the case of a strictly feasible open-loop control (see Dockner et al., 2000, p. 319–320), which implies that we do not allow for individuals to extract all the resources in finite time (see Dockner et al., 2000, p. 321-323 for alternative open-loop controls when this is allowed). Now substituting eq. (89) into eq. (90) and taking the limit yields that 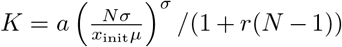. In turn, using the value of this constant in eq. (87) and eq. (89) allows us to determine explicitly the candidate equilibrium control and state paths as

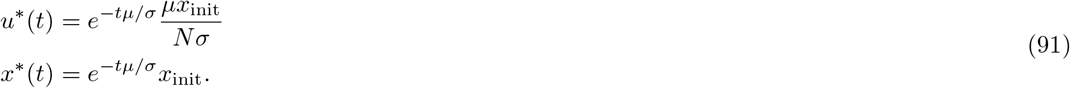

[Note that formally this solution also satisfies open-loop solution with a terminal condition lim_t→∞_ *x*\*(*t*) ≥ 0, which implies a transversality condition given by Note 3 by Sydsaeter et al., 2005, p. 349 and were their *x*_1_ = 0 and λ(*t*) = *e^tμ^*λ*(*t*) > 0 for our problem, which leads together with Note 4 that the terminal condition lim_*t*→∞_ *x*\*(*t*) ≥ 0 simplifies to (90)].

#### 4.2.5 Comparison between closed-loop and open-loop resource extraction rate

Resource extraction crucially depends on relatedness *r* under the (stationary) closed-loop control and resources are extracted much faster when relatedness is low (see Fig. 6). Interestingly, under open-loop control resource extraction is independent of relatedness and corresponds, regardless of population structure, exactly to the open-loop control established for this model in the game theory literature (Dockner et al., 2000, Chapter 12.1), which itself corresponds to the so-called ‘‘cooperative solution” maximizing group payoff (Dockner et al., 2000, Theorem 12.1, p. 320). Hence, relatedness can have no impact on slowing down the extraction rate, which already takes the optimal path from the perspective of the group. Intuitively, one thus expects that the closed-loop equilibrium coincides with the open-loop equilibrium when groups are clonal (*r* = 1), which is indeed the case and again instantiates our broad result that state-feedback has no effect when interacting individuals are clonal.

**Figure 6:**
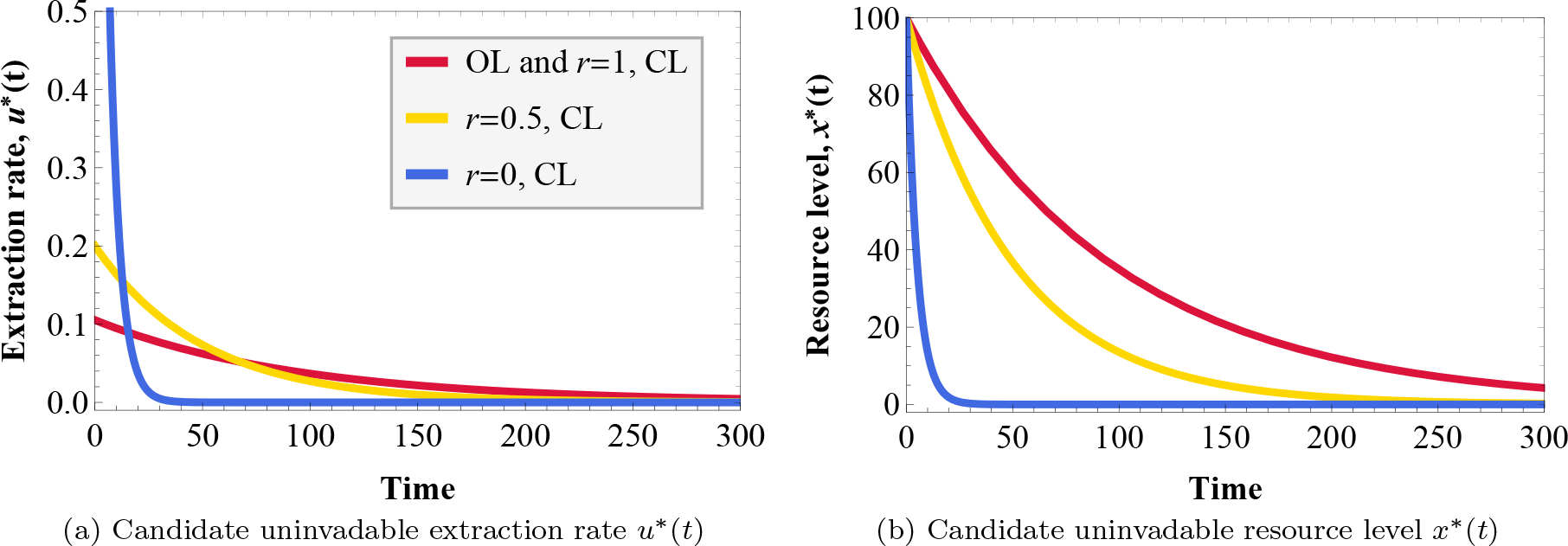
Candidate uninvadable extraction rate (panel a) and resource level (panel b) for closed-loop (CL) and open-loop (OL) traits for different values of average relatedness *r*. Parameter values: *N* = 10, *μ* = 0.01, *σ* = 0.95, *x*_init_ = 100, *T* = 100. Note that if individuals are clones (*r* = 1), the closed-loop and open-loop traits coincide and for open-loop traits extraction rate does not depend on relatedness *r*.

## 5 Discussion

We formalised the directional selection coefficient on a genetically determined function-valued trait when interactions occur between individuals in a group-structured population subject to limited genetic mixing. This selection coefficient describes the directional evolution of a quantitative function-valued trait and determines three relevant evolutionary features. First, it gives the invasion condition of a mutant allele coding for a multidimensional phenotypic deviation (the deviation of a whole function) of small magnitude and takes the form of Hamilton’s marginal rule −*c*+*rb* > 0, where the marginal direct fitness effect –*c* and the marginal indirect fitness effect *b* are given by directional derivatives (formally Gâteaux derivatives). Second, the selection gradient is frequency-independent (same for all allele frequencies) and thus underlies gradual evolution of function-valued traits, since −*c*+*rb* > 0 implies not only initial invasion of the mutant function-valued deviation, but also substitution of the resident ancestral type in the population. Finally, the stationary selection gradient (i.e. when –*c* + *rb* = 0) gives the necessary first-order condition for uninvadability and allows to characterise long-term evolutionary outcomes. While these three features are well known to hold for scalar traits (e.g., Rousset, 2004; Lehmann and Rousset, 2014; Van Cleve, 2015), our derivation of Hamilton’s marginal rule for multidimensional traits generalises them to traits of arbitrary complexity.

Connecting Hamilton’s marginal rule with optimal control and differential game theory, we developed an approach to characterise a necessary first-order condition for the uninvadability of dynamic traits, which applies to both open-loop controls, whose expression is only time-dependent, and closed-loop controls, whose expression is dependent on dynamic state variables as well. We showed that Hamilton’s rule in this context can be decomposed into current inclusive fitness effect, on one side, and future inclusive fitness effect, on the other. The latter effect arises through changes in the states of interacting individuals and depends on the (neutral) shadow values of the states. The shadow value of a state measures how a current change in that state variable affects all future fitness contributions in a population at a demographic equilibrium in the absence of selection, which is given by the neutral current-value reproductive value (or residual fitness). The shadow values are thus central in balancing the trade-off between current and future fitness effects and thus in shaping inter-temporal trade-offs; a feature well-know for open-loop controls (e.g., Perrin and Sibly, 1993 for a review) and that we showed applies equally well to closed-loop traits, which is a result that seems to have neither appeared previously in the differential game theory literature.

Open-loop and closed-loop trait characterisations have sometimes been used in the literature to analyse the same biological phenomena. Our analysis allows for a direct comparison between these two different modes of trait expression. While the selection coefficient takes the same form (Hamilton’s rule) for both, the dynamic constraints are different and this is captured by differences in the dynamics of the shadow value. For open-loop traits, the shadow value dynamics for a given state depends only on how variation in that state variable affects current fitness and state dynamics. For closed-loop traits, the shadow value dynamics depends additionally on the state feedback effect, which captures how a variation in the state variable brings forth variations in traits of all individuals in interaction and this in turn affects current fitness and state dynamics (eq. 35). This causes inter-dependencies between the states of different individuals and inter-temporal effects between trait expressions that are absent under open-loop controls. Analysis of the feedback effect leads to two insights about the role of state-dependence of trait expression in shaping trait evolution. First, the sign of the feedback effect (the sign of eq. 35) determines if the shadow value is larger (for positive feedback effect) or smaller (for negative feedback effect) for closed-loop traits than for open-loop traits. Second, state-dependence of trait expression plays no role if there are no social interactions between individuals or interactions occur only between clones (*r* = 1), in which cases the candidate uninvadable open-loop and closed-loop trait expressions coincide (and the state-feedback term is zero). This means that the use of closed-loop controls appearing in life-history models without interactions between individuals (e.g., Houston et al., 1999) do not lead to different results if instead an open-loop representation of traits would have been used.

We worked out two examples to illustrate these concepts, one of common pool resource production and the other of common pool resource extraction. Interactions under closed-loop trait expression cause individuals to invest less into common pool resource production (and more in extraction) at an evolutionary equilibrium. These results are in line with the literature of economic game theory, where “period commitments” lead to higher levels of cooperation in common pool situations (Meinhardt, 2012). Indeed, an open-loop control can be viewed as a trait (or strategy) committing to its expression over the entire interaction time period, since it cannot be altered in response to some change experienced by individuals. A closed-loop control has no commitment over time since it is expressed conditional on state. Depending on the nature of the interaction between individuals, closed-loop trait expression can then also lead to higher levels of cooperation. A concrete example is an analysis of the repeated prisoner’s dilemma game, where open-loop controls lead to defection, while closed-loop controls can sustain cooperation (e.g., Weber, 2011). The reason why closed-loop strategies are able to sustain cooperation under repeated prisoner’s dilemma is that they allow to condition trait expression on the actions of others, and thus take into account the future threat of punishment. In other words, for closed-loop traits current actions are linked to future ones. Hence, only closed-loop strategies can sustain the reciprocity principle of repeated games by giving rise to incentives that differ fundamentally from those of unconditional trait expression (see Binmore, 2020, p. 87 for a characterisation of this principle).

We finally discuss the scope and limitations of our formalisation. First, concerning scopes, we focused explicitly on the two main types of controls, but also worked out the simplifications that arise under special cases of these controls. In particular, under (closed-loop) stationary controls, where trait expression depends only on state and not on time and the time horizon of the interaction period becomes large, the PDE for the (current-value) reproductive value function no longer depends on time and becomes easier to solve (see “Stationary control result”). Stationary controls have been considered in evolutionary biology and conservation biology, e.g. to model foraging strategies (McNamara et al., 1991; Mangel et al., 1988), web-building behaviour (Venner et al., 2006), adaptive management plans (Chadès et al., 2017). Finally, under (open-loop) constant controls, where trait expression depends neither on state nor on time, the ODEs’ for the shadow values and state dynamics become autonomous, which makes it the simpler case to analyse (see “Constant control result”). Several concrete biological situations fall into this category. For instance, neural networks are dynamical systems whose output is controlled by a finite number of scalar weights (Haykin, 2009), the selection on which is an example of a situation with constant control if weights are taken to be traits evolving genetically (see Ezoe and Iwasa, 1997 for an application to evolutionary biology). Likewise, phenomena as different as gene expression profiles and learning during an individual’s lifespan can be regarded as the outcomes of dynamical systems controlled by a finite number of constant traits (e.g. see respectively Alon, 2020 and Dridi and Akçay, 2018). The case of constant control is thus likely to be widespread in models in evolutionary biology and it would be interesting to work out more explicitly the connection between models for the evolution of learning and those based on control theory. Owing to the presence of both a state dynamics and a decision rule, the control theory approach to trait expression has a universal character (Haykin, 2009, chapter 15), which should thus in principle be able to cover all types of trait expression.

Concerning limitations, we considered deterministic state dynamics but stochastic state dynamics may be interesting to consider in the future by way of applying stochastic optimal control theory (e.g. Kamien and Schwartz, 2012). Perhaps more importantly, we modelled a population reproducing in discrete time, where within each time period individuals can interact for a fixed time interval. As such, the vital rates of individuals can change over this interaction period, but not between interaction periods. Hence, our model with limited dispersal and time-dependent vital rates applies to semelparous species, which covers models with conflict between relatives in annual organisms (Day and Taylor, 2000; Avila et al., 2019). Furthermore, if we allow for complete dispersal between groups (*r* = 0), then our framework can be used to address the evolution of function-valued traits under overlapping generations with time-dependent vital rates as in continuous time classical life history models with and without social interactions (e.g. León, 1976; Schaffer, 1983; Stearns, 1992; Perrin, 1992), but we add the possibility of considering the evolution of closed-loop controls. This scenario is encapsulated in our formalisation because the individual fitness function we used to analyse dynamic trait evolution (eq. 13) takes the same functional form as the basic reproductive number in age-structured populations (and which is sign equivalent to the Malthusian or the geometric growth rate, e.g., Karlin and Taylor, 1981, p. 423-424). As such, our results on closed-loop controls (section 3.2) allow to characterise long-term evolutionary outcomes when the fitness of an individual takes the form of the basic reproductive number. For this situation, our results for open-loop controls (section 3.3) reduce to the standard Pontryagin’s weak principle used in life-history models (e.g. Schaffer, 1983; Stearns, 1992; Perrin, 1992). In order to cover time-dependent vital rates with overlapping generations within groups under limited dispersal, one needs to track the within-group age structure (e.g. Ronce et al., 2000), which calls for an extension of our formalisation. Finally, we did not consider between-generation fluctuations in environmental conditions, which certainly affect the evolution of function-valued traits and it would be interesting to investigate this case. Hence, while our results are not demographically general, our hope is that the present formalisation is nevertheless helpful in providing broad intuition about the nature and conceptualisation of directional selection on phenotypically plastic traits.

## Acknowledgements

We thank Charles Mullon for helpful comments on the manuscript.

## Box 1: Hamilton’s indicators of the force of selection as selection on constant controls

Assume that individual fitness takes the form of the basic reproductive number 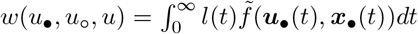, where controls are constant 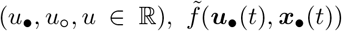 is the fecundity at age *t* and *l*(*t*) is the probability of surviving until age *t* treated as an additional state variable satisfying the dynamics d*l*(*t*)/d*t* = –*μ*(**u**, (*t*), ***x***_•_(*t*))*l*(*t*). Then, it follows from our Constant Control Result that under these assumptions, the selection coefficient on a constant control that acts only at time *t* can be written as

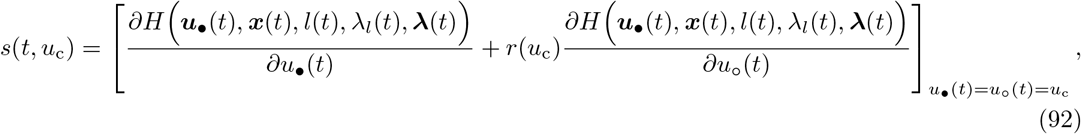

with Hamiltonian

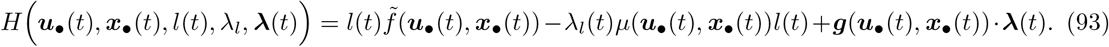

Here, 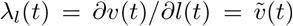 is the shadow value of the cumulative survival probability, which is precisely the neutral current-value reproductive value; namely, Fisher’s reproductive value, since by definition 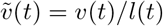 (see also Goodman, 1982; Perrin, 1992 for making this connection and Perrin, 1992; Perrin et al., 1993 for writing the Hamiltonian as in eq. (93) by treating *l*(*t*) as a distinguished state variable).

Assuming no interactions between relatives and that the control affects only fecundity and survival, eq. (92) becomes

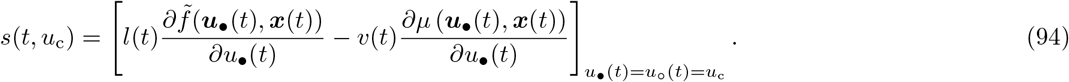

This is consistent with Hamilton’s indicators of the force of selection (Hamilton, 1966). Indeed, for a modifier of fecundity, the first term in eq. (94) is consistent to that of Hamilton’s analysis (e.g., Ronce and Promislow, 2010, eq. 3.3). For a modifier of survival, the second term in eq. (94) is consistent that of Hamilton’s analysis (e.g., Ronce and Promislow, 2010, eq. 3.2, where *μ*(*t*) = – log(*p*(*t*)) and *p*(*t*) is the survival probability at *t* and their generation time *T* is a scaling factor that coverts the first-order perturbation of the basic reproductive number into that of the geometric growth rate, and explains why *T* does not appear in (94) but appears in previous formulations of Hamilton’s indicators of the force of selection on age-specific modifier traits derived from the geometric growth rate of a mutant, e.g. Charlesworth, 1994, chapter 5.1). Finally, we note that in the presence of interaction between relatives, eq. (92) is consistent with results of kin selection in age-structured populations (e.g. Charlesworth, 1994, chapter 5.3.5). Note that the vital rates in eq. (94) can depend in a non-linear way on phenotypic effects and may be time-dependent since they depend on state.

## Appendix A: Derivation of Hamilton’s rule for function-valued traits

In this Appendix, we prove the gradient version of Hamilton’s rule for function-valued traits and show that this provides an invasion implies substitution principle under weak selection (eqs. 4–8). A central concept used in our proof is the notion of a Gâteaux derivative.

### A.1 Gâteaux derivative and point-wise functional derivative

Let 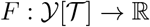 be some functional where 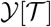 is a vector space over a domain 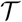 and assume that for some 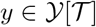 the limit

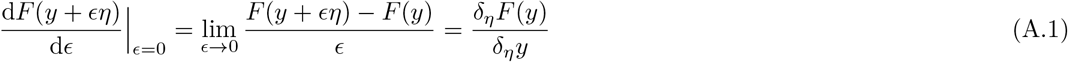

exists for all deviations *η* that satisfy 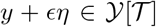 for a sufficiently small non-negative parameter *ϵ*. Then, the function *F* is said to be Gâteaux differentiable at *y*, and *δ_η_F*(*y*)/*δ_η_y* is the shorthand notation for a Gâteaux derivative at *y* in the direction of *η* (Hille and Phillips, 1957, Section 3). The Gâteaux derivative can thus be thought of as a generalization of the directional derivative familiar from finite dimensional spaces. Most rules that hold for ordinary derivatives also hold for Gâteaux derivatives, e.g. Taylor’s theorem and the chain rule (e.g. see Section 2.1C in Berger, 1977, or, Appendix A of Engel and Dreizler, 2013). The Gâteaux derivative can be expressed in terms of point variations (e.g. see Engel and Dreizler, 2013, eq. A.15 and eq. A.28) as

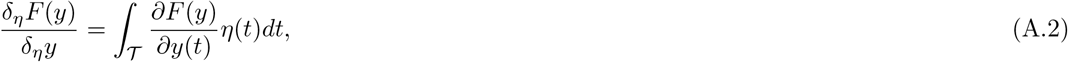

where

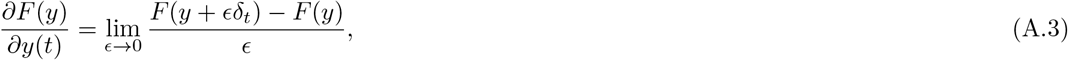

is the point-wise functional derivative of *F* at *y*(*t*) and *δ_t_* is the Dirac measure taking value 1 at *t* and otherwise it is 0. That is, eq. (A.3) is the partial derivative of *F* with respect to *y* at *t* and hence we use the more familiar ‘partial derivative’ notation from finite dimensional spaces. The representation in eqs. (A.2)–(A.3) is useful because it allows, for instance, to take a functional derivative of fitness with respect to the trait, and partition it into a deviation *η*(*t*) and a marginal fitness effect at a specific (single) time point 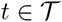, *∂F*(*y*)/*∂y*(*t*) (i.e. a point-wise marginal fitness effect), and only then integrate over the domain 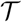.

### A.2 Dynamics of mutant-frequency

Consider that the mutant allele coding for trait *u*_m_ and the resident coding for trait *u*, segregate in the homogeneous island population as described in the main text. Because no individual-level demographic heterogeneity is assumed withing groups (i.e., no class structure), each group can be characterised, from a population genetic state perspective, by the number of mutants that inhabit a given group and we denote the set of all group genetic states with 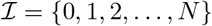. The state of the entire homogeneous island population can thus be described with the vector 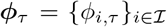 where *ϕ_i,τ_* is the frequency of groups with *i* mutants at demographic time *τ*. Since population size is constant in the homogeneous island population (mean fitness is one), the change in the average frequency Δ*p_τ_* = *p*_*τ* + 1_ – *p_τ_* of the mutant allele from demographic time *τ* to *τ* + 1 (over one life-cycle iteration) can be expressed as

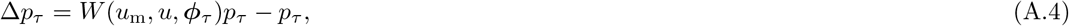

where *W*(*u*_m_, *u, ϕ_τ_*) is the marginal fitness (or lineage fitness) of the mutant allele. Namely, this is the expected number of offspring (including the surviving self) produced by a randomly sampled mutant individual from the collection of all mutants in the population when the distribution of mutants across groups is *ϕ_τ_*. This fitness can be written as the average

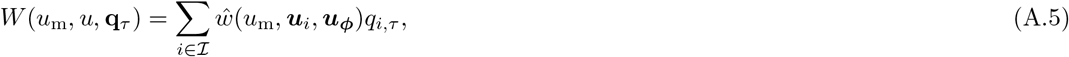

where 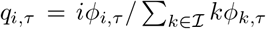 is the probability that a randomly sampled mutant resides in a group with *i* mutants (whence 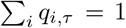) and where ***u**_i_* = (*u*_m_, *u, i* – 1) and ***u**_ϕ_* = (*u*_m_, *u, ϕ_τ_*) are vectors that describe, from the perspective of a mutant sampled in a group with *i* mutants, the distribution of traits among group neighbours (local individuals) and in the groups in the population at large (non-local individuals), respectively. The function 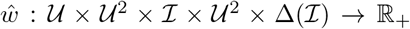 is the individual fitness where 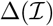 denotes the space of frequency distributions on 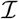 (i.e. the simplex in 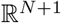), and as such, *ŵ*(*u*_m_, ***u**_i_, **u**_ϕ_*) gives the fitness of a mutant when among its neighbours *i* – 1 individuals have trait *u*_m_ and *N* – (*i* – 1) have trait *u*, and in the groups in the population at large, mutant and resident traits follow the **φ**_τ_ distribution. When the mutant is rare, eq. (A.5) reduces to the invasion fitness of the mutant allele in the homogeneous island population (Mullon et al., 2016, eq. 1).

### A.3 Weak-selection approximation

We now study mutant gene frequency change Δ*p_τ_* assuming small *ϵ*. To that end, it is useful to note that the fitness of a mutant in a group in state *i* can be approximated by writing it in terms of average traits as

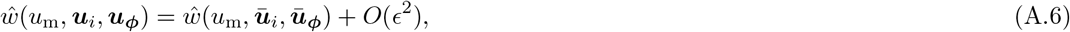

where 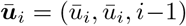 specifies that all group neighbours have the same group average trait 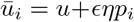 with *p_i_* = [(*i* – 1)/(*N* – 1)] being the frequency of mutants among neighbours, while 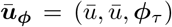 specifies that all non-local individuals have the same average population trait 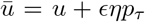 with 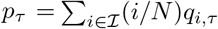 being the average mutant frequency in the population. Eq. (A.6), which has been used for scalar traits (Rousset, 2004, p. 95), follows by Taylor expanding *ŵ*(*u*_m_, ***u**_i_, **u**_ϕ_*) to the first-order about *ϵ* = 0 and using the chain rule (which applies to Gâteaux derivatives, e.g. eq. A.38 in Engel and Dreizler, 2013) to see that the coefficients of the Taylor series involve (at most) Gâteaux derivatives weighted by average allele frequencies. This is an instantiation of the so-called generalised law of mass action (Meszéna et al., 2005; Dercole, 2016) and is secured by the assumption that all individuals within a group that have the same trait are exchangeable (individuals are demographically homogeneous).

Because all non-local (mutant and resident) individuals are considered to have the same average trait (the same is true for group neighbours), 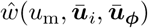 is *de facto* independent of *ϕ_τ_*. This allows us to further simplify the right-hand side of eq. (A.6) by writing

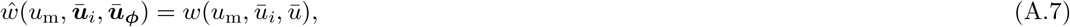

where the function 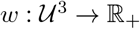 is the (average) fitness function introduced in section 2 of the main text, where we do not need to detail mutant distributions. Hence, *w*(*u*_m_, *ū_i_, ū*) is the fitness of an individual with trait *u*_m_ in terms of only the average trait *ū_i_* of its group-neighbours and the average trait u of individuals in the population.

#### A.3.1 Allele frequency change

Substituting eq. (A.5)–(A.7) into eq. (A.4), we can express the change in allele frequency as

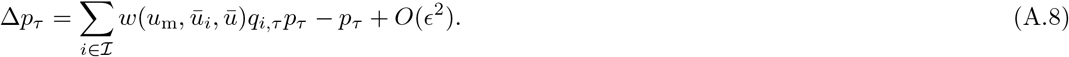

Taylor expanding the fitness function to the first-order about *ϵ* = 0 yields

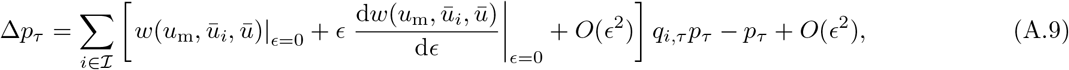

where *w*(*u*_m_, *ū_i_, ū*)|_*ϵ*=0_ = 1 (fitness in a monomorphic population is one), whereby

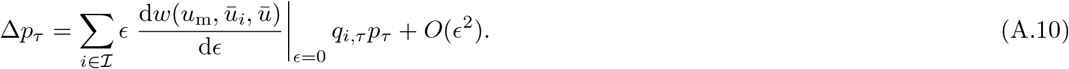

We now apply eq. (A.1) and use the chain rule for Gâteaux derivatives (see e.g. eq. A.38 in Engel and Dreizler, 2013), which produces

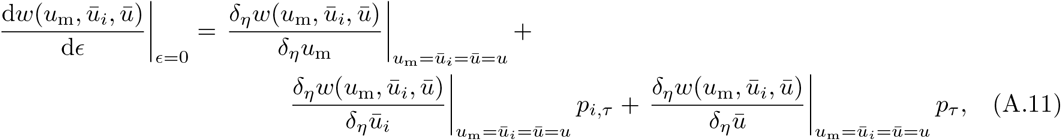

where all partial derivatives here and henceforth are evaluated at the resident value *u*. Since all the partial derivatives are independent of any allele frequency, they give the effects on any individual’s fitness stemming, respectively, from itself, its average neighbour, and an average population member by varying (infinitesimally) trait expression. Hence, the type of the actor is not relevant when evaluating the fitness effects and we can equivalently write eq. (A.11) as

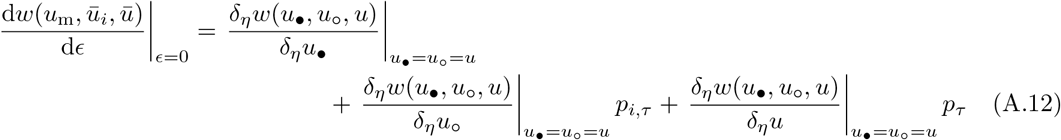

where we took into consideration that the sum of partial derivatives of the fitness function with respect to all of its arguments is zero (since population size is constant, see e.g. Rousset, 2004, p. 96 for scalar traits) and where we replaced the variables *u*_m_, *ū_i_*, and *ū* with *u*_•_, *u*_o_, and *u* (note that we have already substituted the resident trait into the final argument). This will be useful subsequently as it makes clear that fitness effects are independent of individual types and thus allows us to focus attention on the fitness of a focal individual.

Substituting eq. (A.12) into eq. (A.10) gives

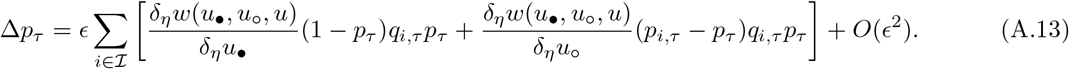

Because 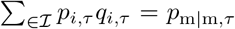 is the probability that, conditional on being a mutant, a randomly sampled neighbour is also a mutant, and *p*_m|m, *τ*_*p_τ_* = *p*_mm, *τ*_ is the probability that two randomly sampled individuals are both mutants (i.e., frequency of mutant pairs), eq. (A.13) can be written

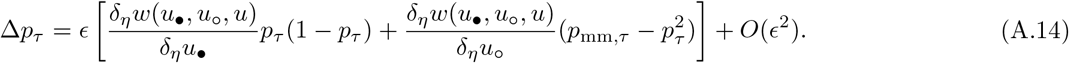

Hence, to the first order in *ϵ*, the dynamics of Δ*p_τ_* is a function of only direct and indirect fitness effects evaluated in the resident population, and the average frequency *p_τ_* and mutant-pair frequency *p*_mm, *τ*_. Further, we only need to study the dynamics of *p*_mm,*τ*_ under neutrality (*ϵ* = 0) because any higher order terms contribute to *O*(*ϵ*^2^) in eq. (A.14). Eq. (A.14) thus generalises to function-valued traits, a standard result for scalar traits (first detailed in Roze and Rousset, 2003 and re-derived a number of times since, e.g., Roze and Rousset, 2004; Rousset, 2004; Roze and Rousset, 2008; Lehmann and Rousset, 2014).

#### A.3.2 Mutant-pair dynamics and relatedness

Using standard population genetic arguments for writing recursions of moments of allelic state (e.g., Jacquard, 1974; Nagylaki, 1992; Roze and Rousset, 2008), we have

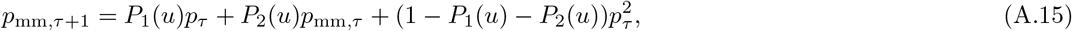

where *P*_1_(*u*) is the the fraction of pairs within groups (of two randomly sampled individuals in the same group without replacement) that descended from the same individual in the previous demographic time step (so that possibly one individual in the pair is the parent of the other in the presence of survival). The quantity *P*_2_(*u*) is the fraction of pairs that have descended from two distinct individuals in the previous demographic time period, and where all these coefficients are constant since they are evaluated under *ϵ* = 0 and thus depend at most on the resident trait *u*. The steady state can be solved explicitly and one gets

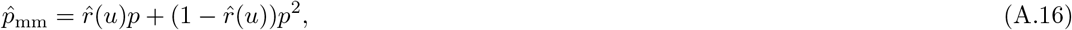

where

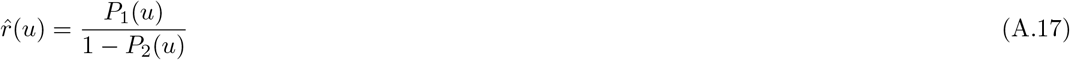

is the relatedness in a patch at the steady state, i.e., the fraction of pairs at the steady state that have a common ancestor in the patch. Owing to neutrality, this is also the probability that a randomly sampled neighbour of a randomly sampled focal individual, carries the same allele as the focal. Moreover, the steady state 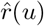 changes continuously with a resident trait whenever *P*_1_(*u*) and *P*_2_(*u*) change continuously.

### A.4 Timescale separation and the invasion implies substitution - principle

We can observe that the dynamics of mutant frequency *p_τ_*, given by eq. (A.14), is dominated by terms of order *O*(*ϵ*), while the mutant-pair frequency *p*_*τ*,mm_, given by eq. (A.15), is dominated by terms of order *O*(1). Hence, when e is small, the variable *p*_mm,*τ*_ undergoes significant fluctuations over the demographic time step Δ*τ* = (*τ* + 1) – *τ* = 1 (one iteration of a life cycle) while *p_τ_* is (nearly) constant. By contrast, *p_τ_* changes significantly over a slower time interval Δ*τ*\* = *ϵ*Δ*τ* while *p*_mm,*τ*_ is near its equilibrium value. We will refer to Δ*τ*^*^ as the evolutionary time step and the phenotypic effect e scales the relationship between evolutionary and demographic time (i.e. one evolutionary time step contains 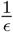 demographic time steps, and equivalently we can write 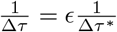).

Combining eq. (A.14) and eq. (A.15), we see that the dynamics of the mutant frequency is thus fully described by the coupled system in demographic time

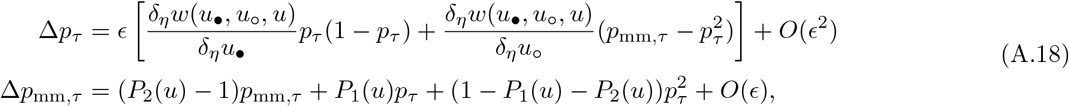

and by a change of variables the system in eq. (A.18) can be equivalently expressed in slow evolutionary time as

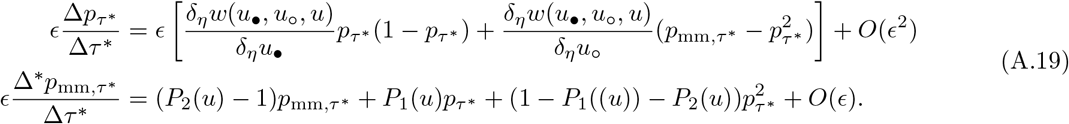

We now separate the demographic and evolutionary timescales (i.e. the timescales of *p*_*τ*,mm_ and *p_τ_*) by letting *ϵ* → 0 and the two last systems above reduce, respectively, to

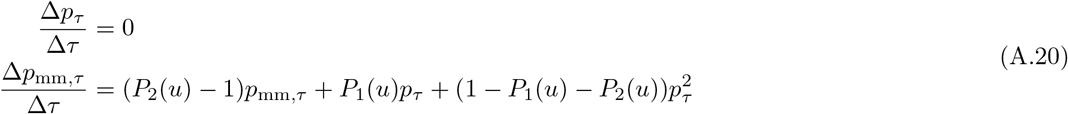

and

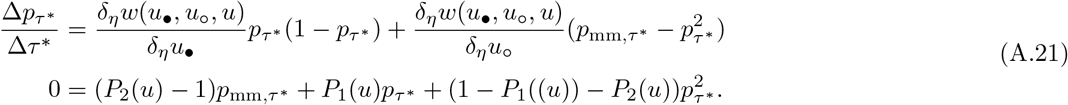

Eq. (A.20) says that in a purely fast demographic time (*ϵ* = 0) the mutant frequency *p_τ_* = *p* stays constant (“frozen in time”), while mutant-pair frequency *p*_*τ*,mm_ changes. Eq. (A.21) says that in a purely slow evolutionary time (*ϵ* = 0) the mutant-pair frequency has reached the steady state 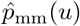 (its solution given in eqs. A.16–A.17), while the mutant frequency *p_τ*_* = *p* changes (thus *p* is in a so-called quasi-steady state – it changes so slowly that it is considered a steady state in one timescale but a fluctuating variable in another). By performing the substitution 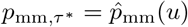 and *p_τ*_* = *p* in eq. (A.21) the dynamics of mutant frequency in slow evolutionary time is

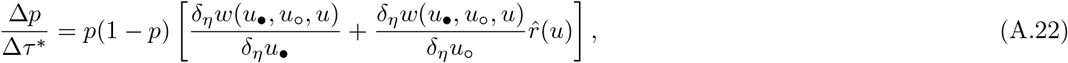

where 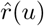 is given in eq. (A.17). Because 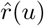 in eq. (A.17) persists under small perturbation of the resident phenotype u (Section A.3.2), we can approximate eq. (A.22) with an equation in fast demographic time whenever *ϵ* is sufficiently small, i.e.

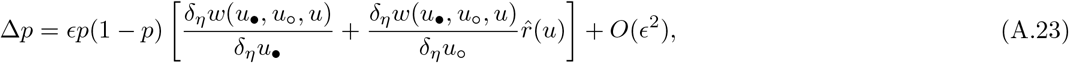

where we used Δ*τ* = 1. This gives us the invasion implies substitution - principle on the time of the demographic process we began with (e.g., eq. A.4). Therefore, we can re-write eq. (A.23) as

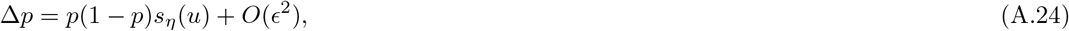

with

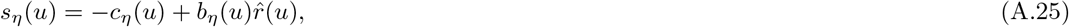

and by using the definition of Gâteaux derivatives in eq. (A.1) we can explicitly write

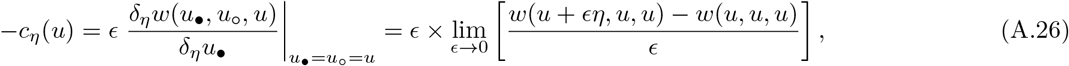

which is the effect a focal individual has on itself if it were to express the mutant phenotype and

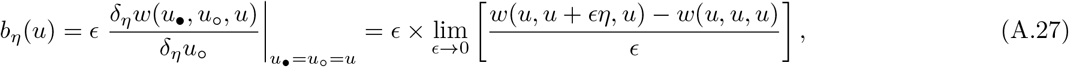

which is the effect that all local individuals have on the focal individual if they were to express the mutant phenotype (where we have likewise substituted u in the second equality). Hence, we have derived eqs. (4)–(5) of the main text.

## Appendix B: First-order condition for state-dependent models

In this Appendix we derive the results of main text section 3. These derivation are based on standard approach of calculus of variations as used in optimal control theory (Liberzon, 2011; Weber, 2011), but our argument will somewhat differ from standard approaches insofar as we will not make use of the Hamilton-Jacobi-Bellman equation, since we are interested only in the necessary first-order conditions (as opposed to necessary conditions in the standard approach). As such, it is important to stress that throughout sections B.1 and B.3, where we derive the dynamics of the (neutral) reproductive value *v*(*t, **x***(*t*); *u*) and the shadow value **λ**(*t*, ***x***(*t*); *u*) = ∇*v*(*t, **x***(*t*); *u*), we evaluate all the traits *u*_•_(*t*) = *u*_o_(*t*) = *u*(*t*) and states *x*_•_(*t*) = *x*_o_(*t*) = *x*(*t*) at some resident values. Only in section B.2 we look at small deviations from the resident population, by analysing the Gâteaux derivatives of the fitness function *w*(*u_•,u__o_, *u*), where we show that we only need to analyse the (neutral) reproductive value *v*(*t*, **x***(*t*); *u*).

For conciseness of notation, we also use the following short-hand notation: for total derivatives w.r.t. time *t* we write 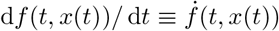, for partial derivatives we write *∂f*(*t, x*(*t*))/*∂x*(*t*) ≡ *f_x_*(*t, x*(*t*)), and second-order partial derivatives we write *∂*^2^*f*(*t, x*(*t*))/*∂x*(*t*)*∂x*(*t*) ≡ *f_xx_*(*t,x*(*t*)). As in the main text, we always use the gradient ∇ notation for gradient with respect to state variables ***x***(*t*).

### B.1 Reproductive value dynamics in a resident population

We here derive the dynamic equations for the reproductive value, eq. (20) of the main text by following the same line of argument as that developed in Metz et al. (2016, see eq. 71), and then we derive an associated equation for the reproductive value that is useful for the other derivations.

#### B.1.1 Partial differential equation for the reproductive value

Recall from eq. (19) of the main text that the reproductive value at time *t* is defined as

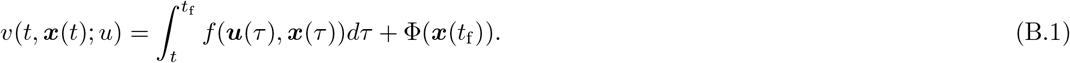

where we recall that the argument ***u*** has been separated with the semicolon in order to emphasise that the controls have been fixed. Hence, for a given ***u*** and initial condition ***x***(*t*) at time t the state trajectory ***x*** is fully determined (i.e. the solution to the ODE in eq. (16) exists and is unique). Because both functions ***u*** and ***x*** are now given functions, the reproductive value in eq. (B.1) is considered to be a function of time t and the initial condition ***x***(*t*) only (strictly speaking it should be a function also of the final time *t*_f_).

In order to derive a dynamic equation of *v*(*t, **x***(*t*); *u*), we consider a very small (but positive) time interval Δ*t* and write eq. (B.1) as

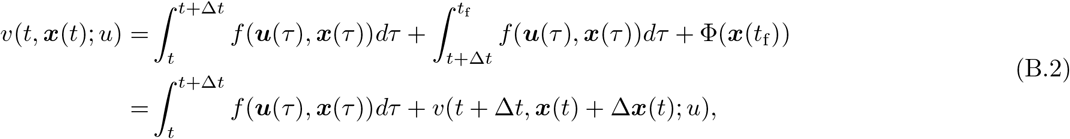

where Δ***x***(*t*) = ***x***(*t* + Δ*t*) – ***x***(*t*) is the change in the state variables over Δ*t* and *v*(*t* + Δ*t, **x***(*t*) + Δ***x***(*t*); *u*) is the reproductive value at *t* + Δ*t* and all arguments have been noted accordingly. Using a first-order Taylor expansion around *t*, we approximate the second term in the second line of eq. (B.2) as

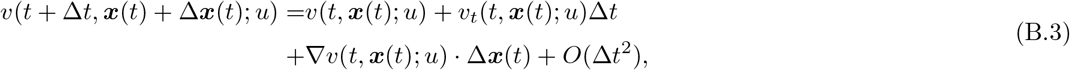

where

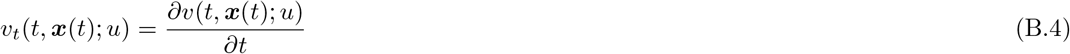

is the partial derivative with respect to the first-argument while

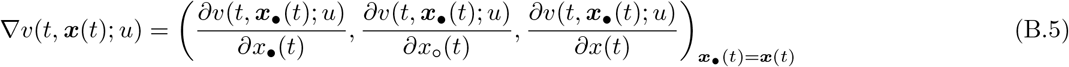

is the vector of partial derivatives with respect to the last argument. Now approximating the first term on the right-hand-side of eq. (B.2) by *f*(***u***(*t*), ***x***(*t*), *t*)Δ*t*, we can write eq. (B.2) as

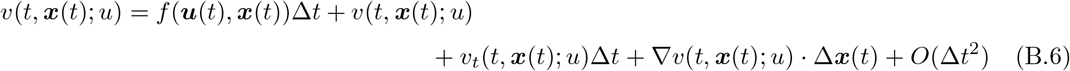

Subtracting *v*(*t, **x***(*t*); *u*) from both sides, dividing by Δ*t*, letting Δ*t* → 0, noting that Δ***x***(*t*)/Δ*t* → ***g***(***u***(*t*), ***x***(*t*)) (as Δ*t* → 0), and rearranging leads to

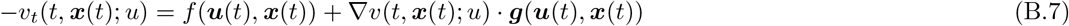

which is a PDE for *v*(*t, **x***(*t*); *u*) with a final condition (f.c.) *v*(*t*_f_, **x**(*t*_f_); *u*) = Φ(***x***(*t*_f_)) for the fixed control path **u**. Eq. (B.7) takes the same form as eq. 71 of (Metz et al., 2016), which was derived under an open-loop control life-history evolution context for a panmictic population and differs with respect to eq. (B.7) in terms of the definition of the arguments.

It is important to stress here that eq. (B.7) is not a form of the so-called Hamilton-Jacobi-Bellman equation for the *value function* evaluated on the optimal control path of optimal control theory (e.g., eq. 3.7 Dockner et al., 2000, chapter 3.2, or eq. 5.10 in Liberzon, 2011 or eq. 3.16 in Weber, 2011), even though it has a similar structure. This is because (i) the reproductive value *v* is here defined to hold for any resident control schedule ***u*** (and is not evaluated at optimality like the value function), and (ii) the value function for our model cannot be computed from the reproductive value of the focal individual, but needs to be computed from the invasion fitness of the mutant, which is the value function in an evolutionary model (invasion fitness is given by eq. A.5 when the mutant becomes rare or eq. 38 in Day and Taylor, 2000, but in the latter case only open-loop traits were allowed).

#### B.1.2 Dynamic equation for the shadow value

Recall that the controls ***u***(*t*) = ***d***(*t, **x***) are functions of ***x***. We now derive the dynamic equation for the shadow value (gradient of reproductive values), which will be useful in later proofs. Taking the gradient of eq. (B.7) with respect to ***x***(*t*), we have

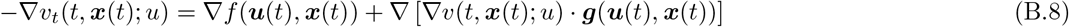

with f.c. ∇*v*(*t*_f_, ***x***(*t*_f_); *u*) = ∇Φ(***x***(*t*_f_)), where

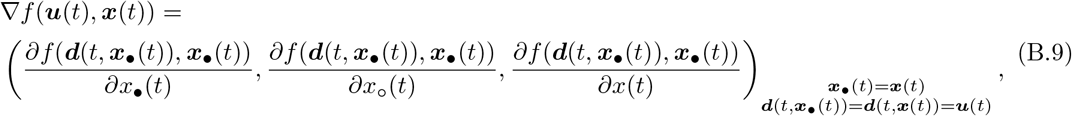

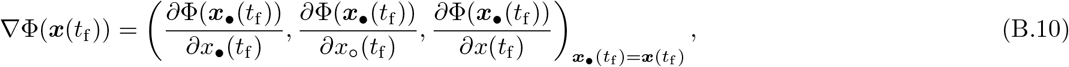

are (column) vectors. Bringing all the terms to the same side and using the chain in rule to expand ∇ [∇*v*(*t, **x***(*t*); *u*) · ***g***(***u***(*t*), ***x***(*t*))], we obtain

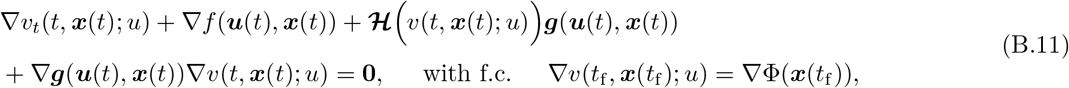

where **0** = (0, 0,0) is a zero (column) vector and

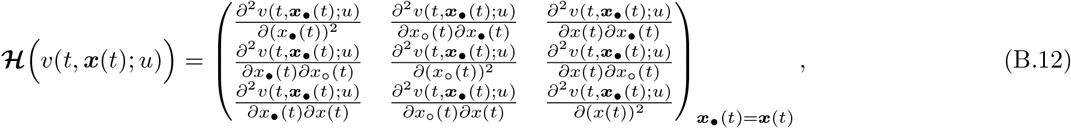

is the Hessian matrix of the reproductive value function and

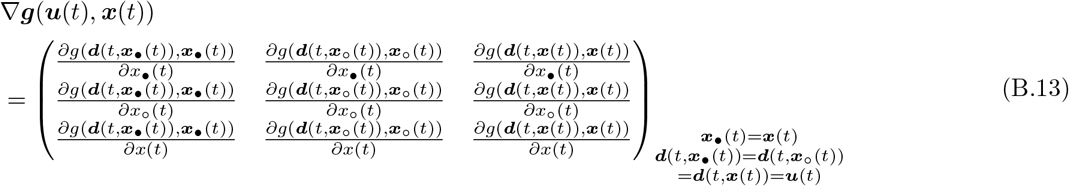

is the gradient of vector ***g***.

Now total differentiating *∂v*(*t, **x***(*t*); *u*) with respect to time and using the property that ***u*** is fixed along a path, we get

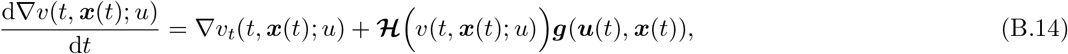

which, on substitution into eq. (B.11), and noting that the order of taking partial derivatives can be changed yields

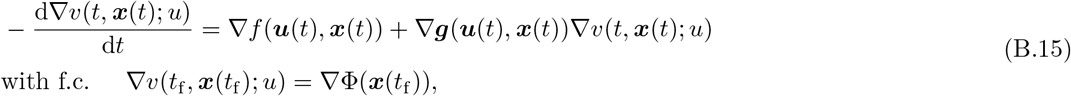

which will be used in the next section.

### B.2 First-order condition and the Hamiltonian

We now turn to deriving the (point-wise) direct effect –*c*(*t, u*(*t*)) and the indirect effect *b*(*t, u*(*t*)), given by eqs. (23) and (24), as well as the point-wise selection gradient for closed-loop traits, eq. (26) and the dynamic equation for the shadow value, eq. (32).

In Appendix A we showed that we can express the direct effect (A.26) and indirect effect (A.27) as Gâteaux derivatives

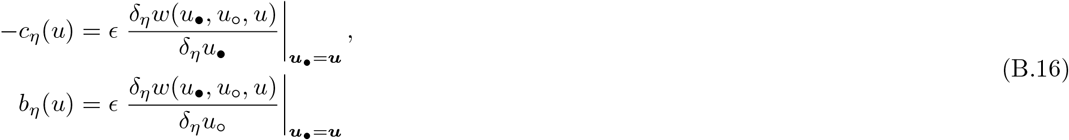

In order to compute these Gâteaux derivatives we first re-write the fitness function *w*(*u_•_, u*_o_, *u*) by augmenting to it a zero quantity containing of adjoint system of constraints (see e.g. Liberzon, 2011, p. 97) and we then we show how to decompose the direct effect –*c_η_*(*u*) and indirect effect *b_η_*(*u*) into point-wise direct effects –*c*(*t, u*(*t*)) and point-wise indirect effects *b*(*t, u*(*t*)), respectively, which allows to characterise the point-wise first-order condition (26).

#### B.2.1 Augmenting the fitness function with an adjoint system of constraints

Recall the individual fitness function eq. (13) of the main text and let us append to it a zero quantity

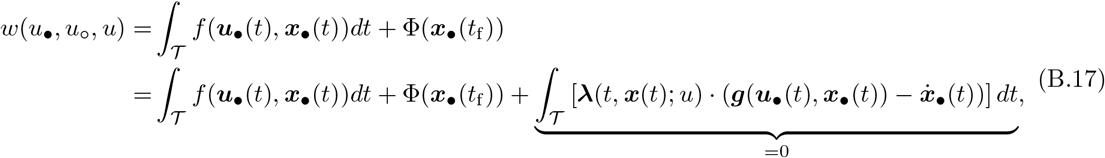

where recalling (eq. 21 of the main text) that **λ**(*t, **x***(*t*); *u*) = ∇*v*(*t, **x***(*t*); *u*) is the shadow value (gradient of reproductive value function). We can integrate the last term in eq. (B.17) by parts

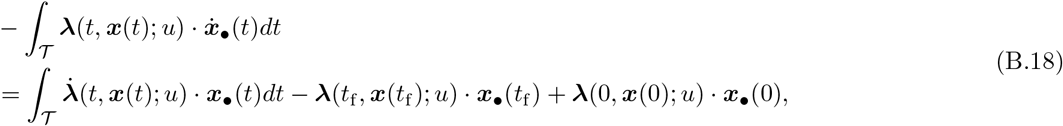

and hence eq. (B.17) becomes

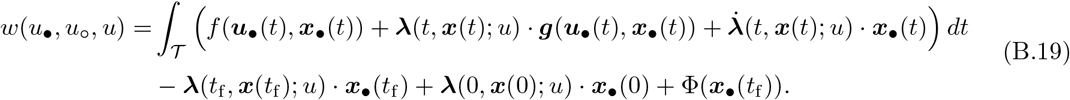

#### B.2.2 Computing the Gâteaux derivatives of the fitness function

We now substitute eq. (B.19) into eq. (B.16) and use eq. (A.2), which yields that we can express the Gâteaux derivatives of the fitness function in terms of point-wise variations as follows

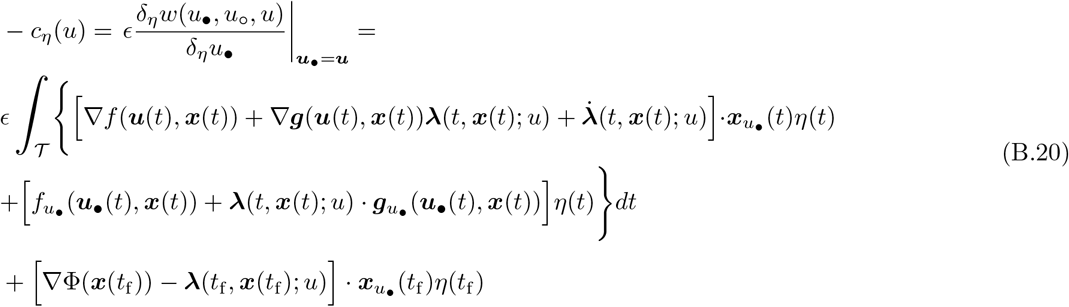

and

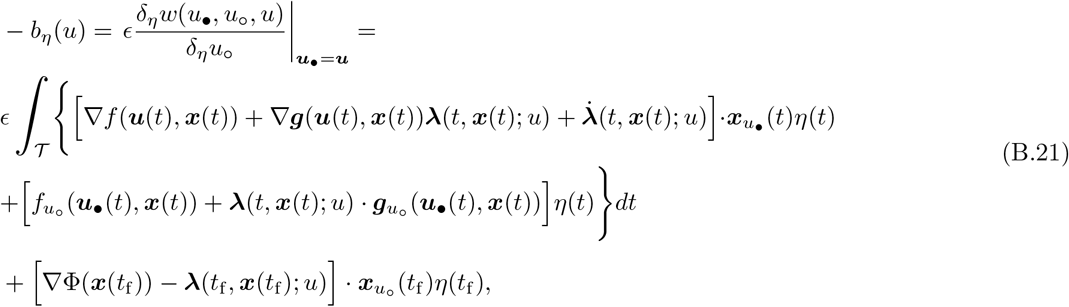

where the term **λ**(0, ***x***(0); *u*) · **x**_•_(0) has disappeared under differentiation because it is a given initial condition, and where all the derivatives under the integrals are evaluated at ***u***_•_(*t*) = ***u***(*t*), ***x***_•_(*t*) = ***x***(*t*) and

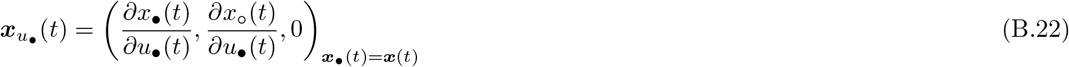

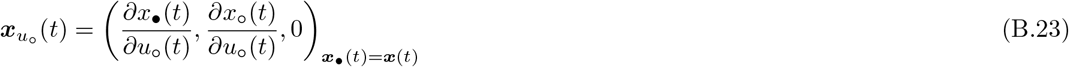

are point variations of ***x***(*t*) caused by variations in *u*_•_(*t*) and *u*_o_(*t*), respectively.

Finally, by definition of the shadow value and from eqs. (B.14) and (B.15) we have

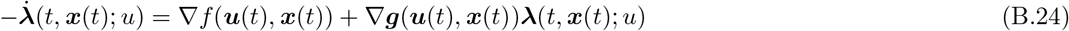

with f.c. **λ**(*t*_f_, ***x***(*t*_f_); *u*) = ∇Φ(***x***(*t*_f_)). Hence, it follows from eqs. (B.14) and (B.15) that the terms (in brackets) multiplying ***x***_*u*_•__, (*t*) and ***x***_*u*_o__(*t*) are zero. Therefore, we can write

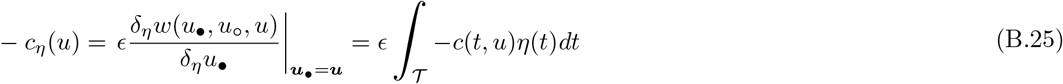

and

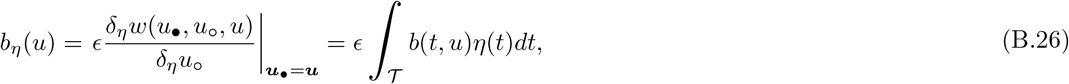

where

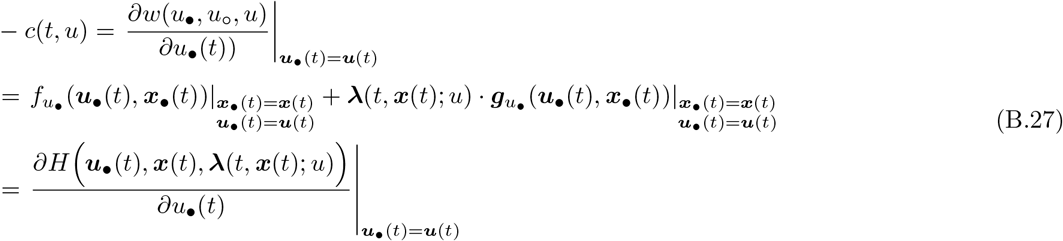

and

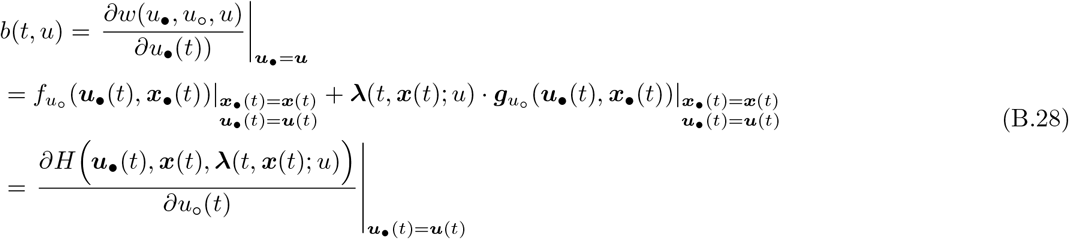

are point variations of *w*(*u*_•_, *u*_o_, *u*) caused by *u*_•_(*t*) and *u*_o_(*t*) (recall eq. A.2) and with this we have derived eqs. (23) and (24) of the main text.

Substituting eqs. (B.27) and (B.28) into eq. (7) of the main text (substituting ***u***\* = ***d***\*(***x***\*), where ***d***\*(***x***\*) = (*d*\*(*t*, *x*\*(*t*)), *d*\*(*t, x*\*(*t*)), *d*\*(*t*, *x*\*(*t*))) and ***x***\* = (*x*,x*,x\**)) and using the definition of the Hamiltonian

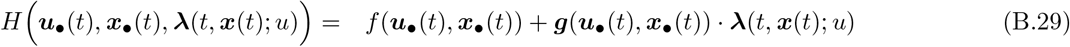

(eq. 25). This yields

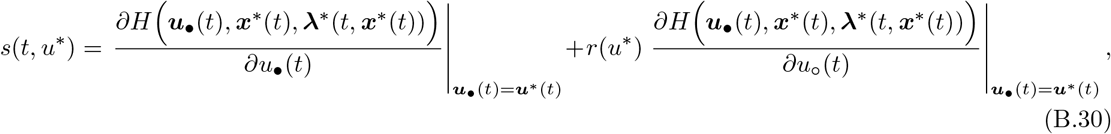

where **λ**\*(*t, **x***\*(*t*)) = ∇*v*(*t, **x***\*(*t*); *u*\*) and the evaluation can be expressed as ***u***\*(*t*) = ***d***\*(*t, **x***\*(*t*)) for closed-loop traits and as ***u***\*(*t*) = ***d***\*(*t*) for open-loop traits. Now recall that the dynamics of *x*\*(*t*) can be obtained from eq. (16) when evaluating it along *u*^*^, which yields

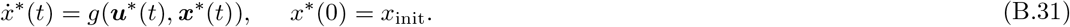

The dynamics of **λ**\*(*t, **x***\*(*t*)) = ∇*v*(*t, **x***\*(*t*); *u*\*) can be obtained from eq. (B.15) and taking into account the Hamiltonian function and evaluating along ***u***\* we obtain for closed-loop control ***u***\* = ***d***\*(***x***\*) path

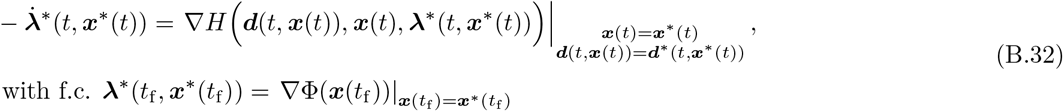

and open-loop control ***u***^*^ = ***d***^*^ path

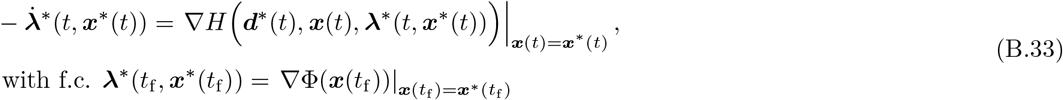

In conclusion, we have derived the point-wise direct and indirect effects, given by eqs. (23) and (24) of the main text (given here by eqs. B.27 and B.28, respectively). In addition we derived the point-wise selection gradient eq. (26) of the main text (here, eq. B.30) along with the dynamic eqs. (32) and (39) on the shadow value function **λ**\*(*t, **x***\*(*t*)) (here, eqs. B.32 and B.33) for closed-loop controls and open-loop controls, respectively. With this we have derived the first-order condition of uninvadability for closed-loop and open-loop controls.

### B.3 Shadow value dynamics and the state feedback

In this section we derive (34)–(33) of the main text; namely, we show that the components of the shadow value dynamics and it depends on higher order derivatives of *v*\*(*t, **x***\*(*t*)). To that end, it will turn out to be useful to explicitly express the control in closed-loop form *u*(*t*) = *d*(*t, **x***(*t*)), unless we are explicitly evaluated at singular path *u*\* = *d*\*(***x***\*). Substituting the Hamiltonian (B.29) into eq. (B.15) yields

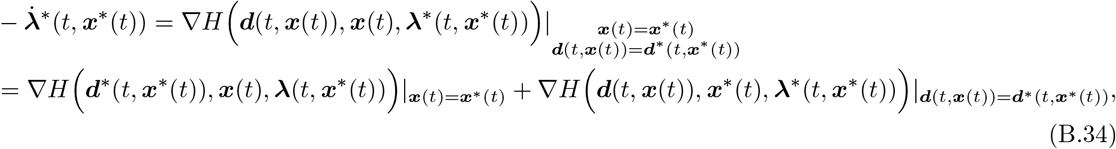

where

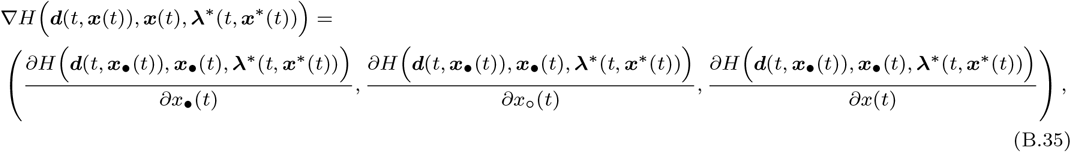

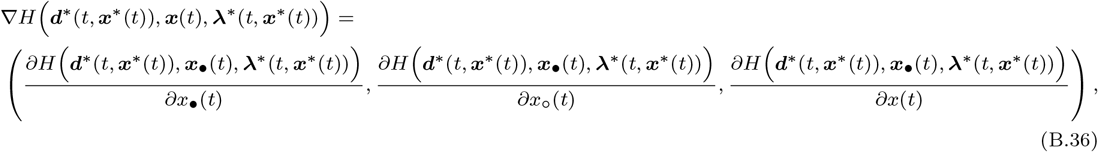

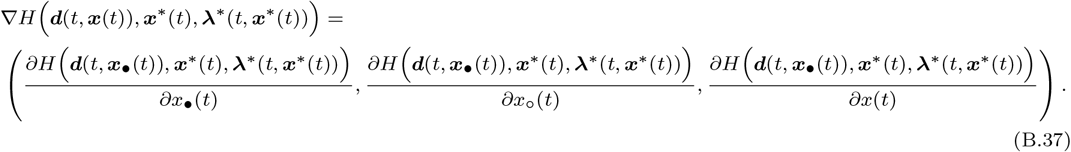

We can express the last gradient (B.37) as

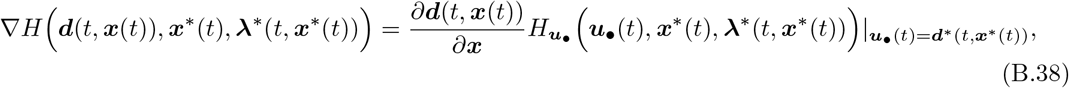

where

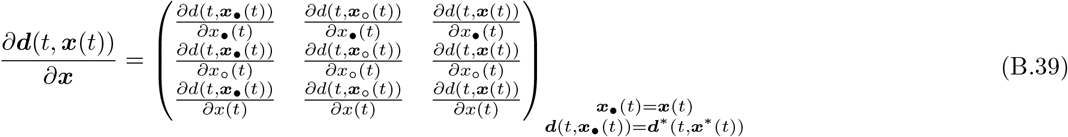

gives all the components of the feedback effect of state variables on trait expressions.

Lets now further explore the elements of a matrix (B.39). From eq. (17) it follows that *∂d*(*t, **x***(*t*))/*∂x*_•_(*t*) = *∂d*(*t, **x***(*t*))/*∂x*_•_(*t*) = 0. From eqs. (17) and (18) it also follows that

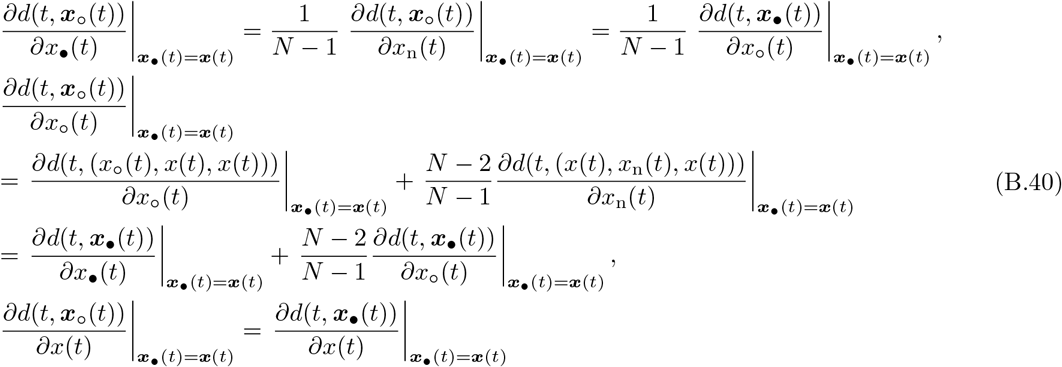

Hence, we can express all the non-zero derivatives in matrix (B.39) as effects of the different actors changing their state on the focal recipient trait expression. Recall the static characterisation (30) from the main text, which holds for interior solutions (when selection gradient (B.30) vanishes)

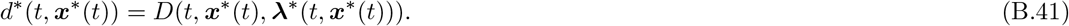

Thus, from eq. (B.41) it follows we can express all the derivations of closed-loop contols *d* in terms of derivations of function *D*, i.e.

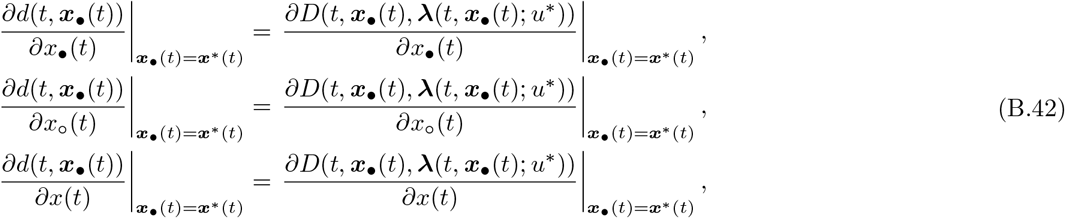

Substituting eqs. (B.40) and (B.42) into eq. (B.39) yields

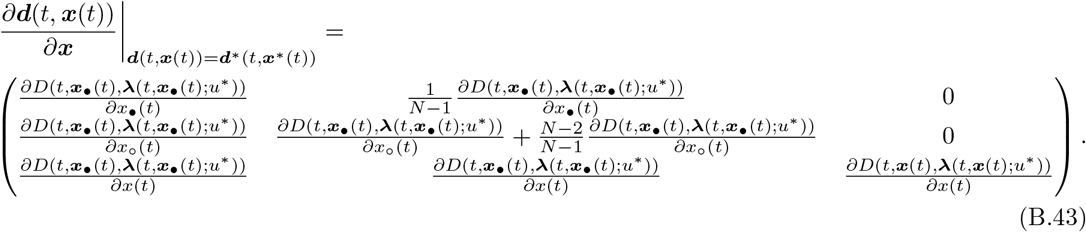

where all the derivatives in the matrix are evaluated at ***x***_•_(*t*) = ***x***(*t*) = ***x***\*(*t*). We can observe from eq. (B.43) that that all the non-zero elements of matrix (B.43) depend on higher-order derivatives of *v*\*(*t, **x***\*(*t*)) and hence eq. (B.34) is not and ODE for **λ**(*t, **x***\*(*t*)) = ∇*v*\*(*t, **x***\*(*t*)).

## Notes

### Competing Interest Statement

The authors have declared no competing interest.

### Summary of Updates

We added Box 1.

